# Long-term herbivore removal experiments reveal different impacts of geese and reindeer on vegetation and ecosystem CO_2_-fluxes in high-Arctic tundra

**DOI:** 10.1101/2023.01.27.525821

**Authors:** Matteo Petit Bon, Brage B. Hansen, Maarten J. J. E. Loonen, Alessandro Petraglia, Kari Anne Bråthen, Hanna Böhner, Kate Layton-Matthews, Karen H. Beard, Mathilde Le Moullec, Ingibjörg S. Jónsdóttir, René van der Wal

## Abstract

1. Given the current and anticipated rates of global change, with associated shifts in herbivore population densities, understanding the role of different herbivores in shaping ecosystem structure and processes is critical for predicting ecosystem responses. Here, we examined the controls exerted by migratory geese and resident, non-migratory ungulates, two dominating yet functionally contrasting herbivores, on the rapidly warming Arctic tundra.
2. We collected vegetation and ecosystem carbon flux data at peak plant growing season in the two longest running herbivore removal experiments in high-Arctic Svalbard. Herbivore exclosures had been set up independently in a wet habitat utilised by barnacle geese (*Branta leucopsis*) in summer and in mesic-to-dry habitats utilised by wild reindeer (*Rangifer tarandus platyrhynchus*) year-round.
3. Excluding geese produced vegetation state transitions from heavily grazed, moss-dominated (4 g m^-2^ dry weight of live aboveground vascular plants) to ungrazed, graminoid-dominated (60 g m^-2^; after 4-yr exclusion) and then horsetail-dominated (150 g m^-2^; after 15-yr exclusion) tundra. This caused large increases in vegetation carbon and nitrogen pools, dead biomass and moss-layer depth. Modifications in nitrogen concentrations and carbon-to-nitrogen ratios of vegetation and soil suggested overall slower nutrient cycling rates in the short-term absence of geese. Long-term goose removal quadrupled the net ecosystem carbon sequestration by increasing gross ecosystem photosynthesis more than ecosystem respiration.
4. Excluding reindeer for 21 years also produced detectable, but weaker, increases in live and dead biomass, vegetation carbon and nitrogen pools, moss-layer depth and ecosystem respiration. Yet, reindeer removal did not alter the chemistry of either vegetation or soil, nor net ecosystem carbon sequestration.
5. Our findings suggest that, though both herbivores were key drivers of ecosystem structure and processes, localised effects of geese, highly concentrated in space and time, are larger than those exerted by more widely dispersed reindeer. We illustrate that the impacts of herbivory across the tundra landscape are contingent on the habitat utilised for foraging, its sensitivity, the exerted grazing pressure, and herbivore characteristics. Our results underscore the conspicuous heterogeneity in how Arctic herbivores control ecosystem functioning, with important implications under current and future global change.

## 1 INTRODUCTION

Vertebrate herbivores are found in most terrestrial ecosystems, and their key role in shaping vegetation and ecosystem structure and function has been recognised for decades in systems as diverse as savannas (McNaughton 1985), temperate grasslands (Frank & Groffman 1998), boreal forests (Pastor & Naiman 1992) and the Arctic tundra (Jefferies et al., 1994). Due to a mixture of anthropogenic factors (Van Eerden et al., 2005), densities of herbivore populations are undergoing drastic changes throughout the Arctic (Uboni et al., 2016; Fox & Madsen 2017). Thus, with the Arctic now experiencing the fastest rate of climate warming on Earth (IPCC 2021), together with the potential for herbivory to at least partly modulate climate effects on vegetation and ecosystem processes (Sjögersten et al., 2008; Olofsson et al., 2009; Leffler et al., 2019), research on how herbivores affect tundra ecosystems has intensified (Barrio et al., 2016). This growing body of studies has not only been informative about the effects of herbivores on vegetation and ecosystem functioning (Beard, Kelsey, et al., 2019), but also showcased that herbivore impacts are far from homogeneous and often depend on the species involved, as well as the ecological context (Sjögersten et al., 2008; Sundqvist et al., 2019; Petit Bon, Inga, et al., 2020). Yet, these insights have mainly emerged when synthesising outcomes from different studies, generally conducted in various Arctic systems and in different years (Forbes et al., 2019; Soininen et al., 2021). Herein, by sampling two independent sets of long-term herbivore exclosures found in the same Arctic system, we illustrate how geese and *Rangifer* (reindeer/caribou), two circumpolar-distributed herbivores, vary in their impacts on tundra vegetation and ecosystem carbon (C) exchange rates.

Herbivores influence standing biomass and composition of tundra vegetation via direct and indirect mechanisms. By directly consuming plant tissue, herbivores generally reduce plant biomass, though no effects or even biomass increases have been reported (Wegener & Odasz-Albrigtsen 1997; Bråthen & Odasz-Albrigtsen 2000). This is because by shortcutting the litter decomposition pathway, through the direct deposition of soluble nutrients (mainly nitrogen [N]) in the form of animal excreta (Van der Wal et al., 2004), and by reducing the depth of the moss-layer through trampling (Tuomi et al., 2021), indirectly increasing soil temperatures (Van der Wal & Brooker 2004; Gornall et al., 2009), herbivores can hasten nutrient cycling rates and enhance primary productivity. This is how Arctic herbivores can promote and perpetuate a graminoid-dominated vegetation state in the tundra (Van der Wal 2006; Ravolainen et al., 2020), whereby they also modify the quality (N concentration and C-to-N ratio [C:N]) of preferred forage plants (Olofsson et al., 2001; Petit Bon et al., 2022). At the extreme, however, if herbivore pressure exceeds certain thresholds, resulting alterations to the plant-soil system can spark positive feedback loops that lead to the formation of near-irreversible, degraded vegetation states (Jefferies et al., 2006).

Through their influence on vegetation and soil, Arctic herbivores can alter C dioxide (CO_2_) emissions from the tundra. A herbivore-induced decrease in plant biomass can directly reduce the potential for C uptake of the ecosystem (Van der Wal et al., 2007; Sjögersten et al., 2008; Metcalfe & Olofsson 2015). Yet, reduced biomass inherently translates to lower ecosystem respiration, potentially levelling out the loss of C at the ecosystem level. Further, herbivores can indirectly enhance C emission by stimulating soil respiration in response to reduced moss-layer depth and hence higher soil temperatures (Gornall et al., 2009). However, this may not be observed if the net effect of herbivory is that of either rising primary productivity by accelerating nutrient cycling rates (Ylänne & Stark 2019) or decreasing soil respiration by limiting the amount of C available for microbes through reduced plant biomass (Stark & Grellmann 2002). Understanding how herbivores affect tundra CO_2_-fluxes may thus provide key information to predict the role of herbivory in the C balance of Arctic regions. This is critical considering that approximately a third of the World’s soil C is stored in permafrost regions (Tarnocai et al., 2009), with large potential for positive feedback to the climate system if released (Schuur et al., 2015).

Arctic ecosystems host a range of different vertebrate herbivores (Speed et al., 2019); how these herbivores impact ecosystem functioning might vary between taxonomic and functional groups, and their pattern of resource utilization in space and time. For example, spatially co-occurring mammalian herbivores with contrasting size and resource utilisation patterns, such as small rodents and reindeer, have been shown to differentially affect tundra vegetation (Ravolainen et al., 2011; Petit Bon, Inga, et al., 2020) and ecosystem CO_2_-fluxes (Metcalfe & Olofsson 2015). Similarly, herbivores that co-occur less in space might differentially alter ecosystem functioning in their respective habitats (Jefferies et al., 1994). Migratory geese are well known for exerting a strong pressure on the Arctic tundra during restricted periods of the year, i.e., following arrival in spring and while rearing their young throughout the summer (Gauthier et al., 1995). Moreover, the intensity of goose herbivory can be variable, with wet habitats experiencing much higher levels of grazing than drier habitats (Madsen et al., 2011), indicating that the effect of geese on tundra ecosystems is likely to be spatially concentrated (Speed et al., 2009). Contrastingly, reindeer graze and browse tundra vegetation year-round, often ranging widely and utilising both wet and drier habitats (Hansen et al., 2010). Furthermore, their highly dispersed use of resources (Iversen et al., 2014) suggests that reindeer are likely to affect tundra ecosystems more evenly in space, and across larger parts of the landscape, compared to geese.

Due to the paucity of long-term herbivore removal experiments in the Arctic, and even more so in the high-Arctic (Forbes et al., 2019), studies on how contrasting herbivores influence various tundra ecosystems in Arctic landscapes are scarce. With the Arctic rapidly changing, there is an urgent need for simultaneous experiments that use comparable designs to help us understand the role of different herbivores for tundra ecosystem functioning. In this study, we quantify grazing impacts of barnacle geese and wild Svalbard reindeer, which populations in the high-Arctic Svalbard have increased considerably in size and spatial extent during the last decades (Fox et al., 2010; Le Moullec et al., 2019). We took advantage of two running long-term herbivore removal experiments, one set up in a wet habitat heavily utilised by the post-breeding and moulting migratory geese and one in mesic-to-dry habitats largely utilised by the resident, non-migratory reindeer. Our goal was thus to quantify how tundra vegetation and associated ecosystem processes respond to the long-term removal of either the respective herbivore from its key habitat. We focused on herbivore-induced alterations in (i) plant biomass and vegetation composition, (ii) concentrations and pools of C and N in vegetation, and (iii) instantaneous ecosystem CO_2_-fluxes. Changes in standing-dead biomass and moss-layer depth, as well as in soil C and N concentrations, were also quantified to draw a more complete picture of the extent to which these functionally different herbivores may influence the high-Arctic tundra.

## 2 MATERIAL AND METHODS

### 2.1 Study systems

This study utilised the two longest running, fully replicated herbivore removal experiments found in the archipelago of Svalbard (Norway), in the European high-Arctic. The two study sites (near the settlement of Ny-Ålesund and in the valley of Semmeldalen) are about 130 km apart, yet both are located on the largest island of the archipelago, Spitsbergen (Figure 1a). The Ny-Ålesund site is situated on the shore of the bay of Kongsfjorden (Thiisbukta), whereas the Semmeldalen site is situated in the inner part of the ca. 8-km long, U-shaped, Semmeldalen valley in the central part of Spitsbergen. Characteristics of the study sites are given in Table 1a.

**TABLE 1.** Characteristics of the study sites.

|  |  | **Ny-Ålesund** | **Semmeldalen** |
| --- | --- | --- | --- |
| **(a)** | Location | 78° 55' N, 11° 55' E | 77° 59' N, 15° 22' E |
|  | Altitude | 5-10 m a.s.l. | 150-165 m a.s.l. |
|  | Mean annual temperature | -4.3 [-2.1] °C | -4.0 [-1.8] °C |
|  | Mean July temperature | 5.6 [6.2] °C | 6.8 [7.2] °C |
|  | Annual precipitation | 465 [749] mm | 204 [293] mm |
| **(b)** | Main herbivore habitat utilisation | Geese (June-July) | Reindeer (year-round) |
| | Trend in main herbivore population | $\lambda = 1.02$ (geese) <sup>(a)</sup> | $\lambda = 1.09$ (reindeer) <sup>(b)</sup> |
|  | Mean main herbivore metabolic biomass in summer (June-July) <sup>(c)</sup> | 560 Kg geese km<sup>-2</sup> | 154 Kg reindeer km<sup>-2</sup> |
|  | Mean main herbivore metabolic biomass throughout the year <sup>(d)</sup> | 93 Kg geese km<sup>-2</sup> | 154 Kg reindeer km<sup>-2</sup> |

**(a)** Climate data are for the 30-year period 1989-2018 from the climate station in Ny-Ålesund (for Ny-Ålesund site) and from the climate station at Svalbard airport (for Semmeldalen site), which is located at 28 m a.s.l., ~30 km northwest of the study site [in brackets, data for the sampling year 2018] (www.met.no). **(b)** Estimated mean annual population growth rate ( $\lambda$ ) for <sup>(a)</sup> barnacle geese for the period 2004–2017 (data from Layton-Matthews et al., 2019) and <sup>(b)</sup> reindeer for the period 1998-2018 (data from Solberg et al., 2022). Estimated metabolic biomass of geese (Ny-Ålesund, period 2004-2017; mean population size $N = 693$ , mean body mass = 1.6 kg, area = 2 km<sup>2</sup>) and reindeer (Semmeldalen, period 1998-2018; mean population size $N = 898$ , mean body mass = 52 kg, area = 300 km<sup>2</sup>), separately for <sup>(c)</sup> summer only and <sup>(d)</sup> whole year (data from references above).

**FIGURE 1.**
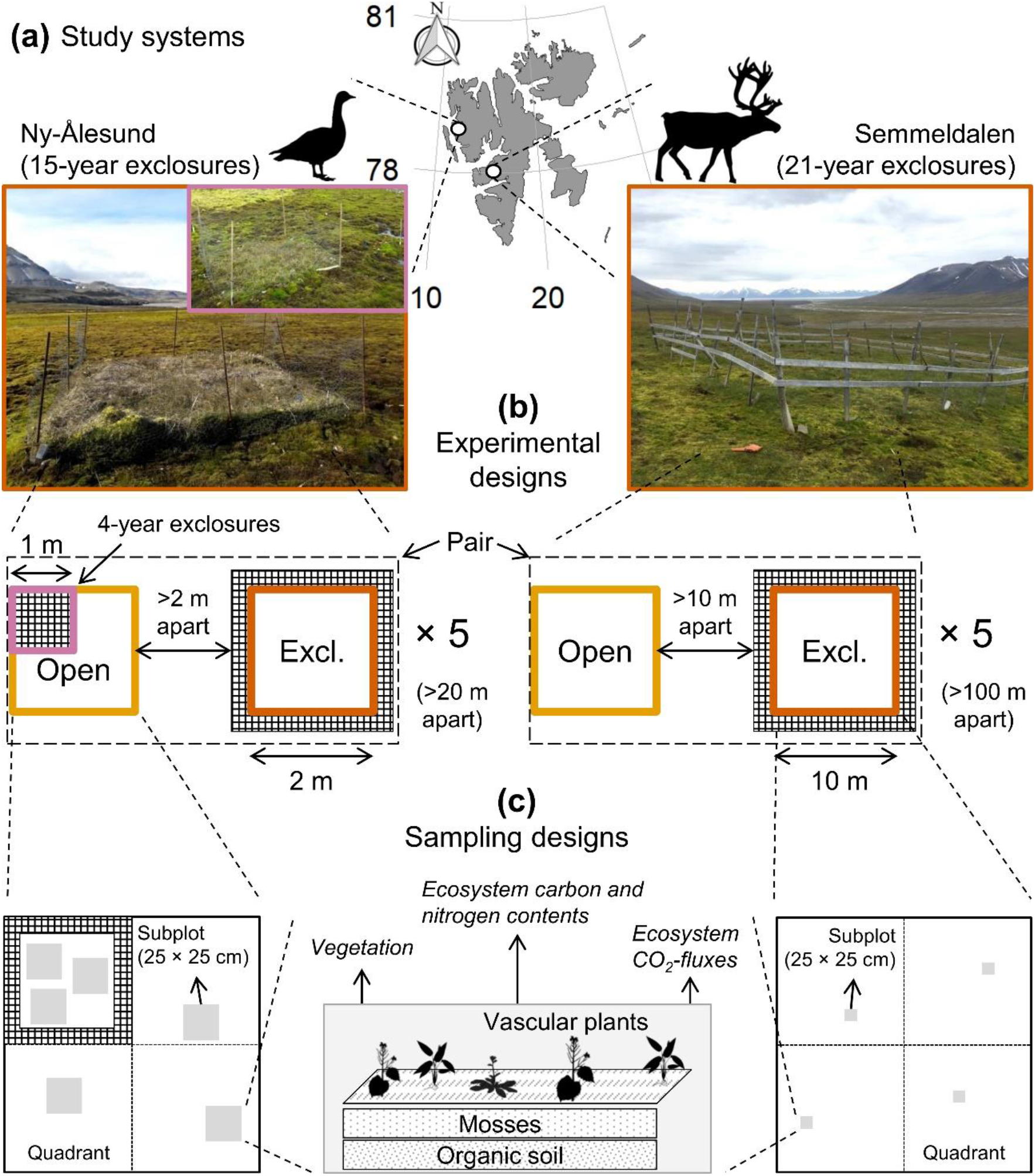
Schematic of the experiments at the two study sites. **(a)** Study systems, **(b)** details on the spatial structure of the experimental designs (main pictures show examples of the long-term exclosures at the two experimental sites in the sampling year 2018; the inset for Ny-Ålesund shows a short-term exclosure), and **(c)** sampling designs adopted for data collection (in total, 45 and 40 subplots have been sampled in Ny-Ålesund and Semmeldalen, respectively). The drawing in **(c)** summarises the focal ecosystem properties considered in this study, which are outlined in the corresponding sections throughout the Article.

There are only two resident (i.e. year-round) vertebrate herbivores in Svalbard, the wild Svalbard reindeer (*Rangifer tarandus platyrhynchus*) and the Svalbard rock ptarmigan (*Lagopus muta hyperborea*). In summer, however, several species of migratory birds breed in Svalbard, among which two geese, the barnacle goose (*Branta leucopsis*) and the pink-footed goose (*Anser brachyrhynchus*), have become exceptionally abundant (Fox & Madsen 2017) and extensively utilise terrestrial wet habitats across the archipelago.

Although the two long-term experiments were independently set up to better understand the influence exerted of either migratory barnacle geese (Ny-Ålesund) or resident, non-migratory reindeer (Semmeldalen) on Svalbard’s ecosystems, they also provide a unique opportunity to address how these herbivores differentially affect the habitat they most extensively utilised across the tundra landscape.

The Ny-Ålesund area has borne an increasing population of barnacle geese for approximately 40 years (Table 1b; Layton‐Matthews et al., 2019), which heavily utilise wet habitats (our focus here, see below). Reindeer are now also abundant on Brøggerhalvøya (the peninsula where Ny-Ålesund is located), but they infrequently use wet habitats in and around Ny-Ålesund and are more commonly observed grazing within drier habitats (MJJEL and MPB, *personal observation*), and particularly so outside the growing season. This renders our study site in Ny-Ålesund a representative example of a plant-herbivore system that is intensively grazed by geese, but only for a limited period of 3-to-5 weeks when vascular plant growth is the greatest. During this period, adult geese undergo full wing moult and goslings grow up, conditions which limit their spatial range. By the time adults and young geese can fly, they leave the area.

In contrast, the Semmeldalen study site is grazed year-round, predominantly by reindeer, a “mixed grazer/browser” herbivore that is less spatially restricted (home-range size of a few km^2^; Tyler & Øritsland 1989) and with a much wider utilisation of plant species in their diet (Hiltunen et al., 2022). The reindeer population in the study area, which has tripled in size during the last 20 years (Table 1b; Solberg et al., 2022), extensively utilises mesic-to-dry habitats (our focus here, see below). Pink-footed geese have also increased in numbers within the valley, as in the rest of the archipelago (Madsen et al., 2017), but they are generally confined to wet habitats (Speed et al., 2009), and only sporadic signs of goose activity were observed at the exclosure location (RvdW and MPB, *personal observation*).

### 2.2 Experimental sites and setup

Detailed experimental site and study design descriptions are given elsewhere (Ny-Ålesund: Sjögersten et al., 2011; Semmeldalen: Van der Wal & Brooker 2004), but a summary is provided here. Plant names follow the Svalbard Flora (https://www.svalbardflora.no) and Frisvoll and Elvebakk (1996).

The Ny-Ålesund experimental site was established in 2003 in a wet habitat, where the growth of the goose population had already strongly impacted the vegetation (Van der Wal & Hessen 2009) through suppressing grasses (mainly *Poa arctica*), forbs (e.g. *Saxifraga* spp. and *Cardamine* spp.) and the horsetail *Equisetum arvense. Calliergon* spp. are the dominant mosses. Five pairs of exclosure and open-grazed plots were set up within a 1-km^2^ area (Figure 1b). Exclosures were built with 50-cm high poultry netting with metal strings across the top to prevent geese from entering. Five additional exclosures were built in 2015 by fencing off a randomly selected corner (quarter) of each open-grazed plot.

The Semmeldalen experimental site was established in 1997 in two different, but moss-dominated, habitats: mesic grass-rich meadow and dry *Luzula*-heath. Across habitats, common vascular plant species are the grasses *P. arctica, Festuca rubra richardsonii* and *Alopecurus ovatus;* the rush *Luzula confusa;* the forbs *Bistorta vivipara* and *Stellaria longipes;* the deciduous dwarf-shrub *Salix polaris;* and the horsetail *E. arvense*. Dominant mosses are *Sanionia uncinata, Tomentypnum nitens* and *Aulacomnium* spp. Five pairs of exclosure and open-grazed plots were established within a 10-km^2^ area (Figure 1b). Three pairs were set up in mesic grass-rich meadows and two pairs in dry *Luzula*-heaths. Exclosures are 150-cm high and made of timber.

We did not observe signs of herbivore activity inside the exclosures during the sampling season of 2018. Yet, the presence of an old reindeer winter-pellet in one of the exclosures in Semmeldalen suggested that reindeer may have accessed these plots over the years, though the pellet might have been transported by the wind on the hard snow surface. Regardless, reindeer access is most likely to happen in winter when the snow can sometimes level out with fences and allows reindeer to step over; however, no over-winter cratering (Beumer et al., 2017) took place. Hence, we consider exclosures to effectively remove the herbivory pressure on vegetation and underlying soil system.

### 2.3 Sample collection and processing

#### 2.3.1 Sampling design

Around the peak of the growing season in 2018 (Ny-Ålesund: 15-21 July; Semmeldalen: 25-31 July), an intense sampling campaign was undertaken to determine how Svalbard’s tundra ecosystems were responding to the long-term removal of geese (Ny-Ålesund; 15-yr exclusion) and reindeer (Semmeldalen; 21-yr exclusion). In Ny-Ålesund, short-term exclosures (4-yr-old) were also sampled to help interpret long-term responses of the wet habitat to goose removal.

To account for some of the spatial variation within plots, measurements of focal ecosystem properties (vegetation, ecosystem C and N contents and CO_2_-fluxes; see below) were taken in three (Ny-Ålesund) and four (Semmeldalen) subplots at each plot (Figure 1c). Plots were divided in quadrants and one subplot was determined at random within each quadrant. No such quadrant randomisation took place within the 4-yr-old exclosures in Ny-Ålesund due to their limited size.

#### 2.3.2 Vegetation: vascular plants and mosses

We assessed aboveground vascular plant biomass using the point intercept frequency methodology (PIM -Bråthen & Hagberg 2004). PIM was performed at each subplot using a sampling frame with 15 points. A pin (3 mm diameter) was lowered from above onto the moss or soil at each point and the number of contacts (intercepts) between the pin and each live vascular plant species was counted. Standing-dead graminoid material was also a possible intercept and recorded separately. We converted the number of intercepts for each species into plant biomass (g m^-2^, as dry weight [dw]) following Bråthen and Hagberg (2004) and by using correlation coefficients in Petit Bon et al., (2021). We grouped vascular plant species into five broadly-classified plant functional types (PFTs), namely forbs, grasses, horsetails, rushes and deciduous dwarf-shrubs (species data is given in Appendix 1, Table S1 of the Supporting Information).

We measured moss-layer depth (separately for the photosynthetically-active green part and the nearly-decomposed brown part) in two randomly selected, yet opposing, corners of each subplot.

#### 2.3.3 Ecosystem C and N contents

We assessed leaf C and N concentrations (%dw) in five focal vascular plant species belonging to the five PFTs outlined above: *B. vivipara* (forb), *P. arctica* (grass), *E. arvense* (horsetail), *L. confusa* (rush) and *S. polaris* (deciduous dwarf-shrub). Combined, these species made up on average 84% and 92% of the aboveground vascular plant biomass at the Ny-Ålesund and Semmeldalen sites, respectively. Moreover, we quantified their total leaf C and N pools (g m^-2^ dw). Finally, we assessed C and N concentrations of mosses and soil.

Vascular plant leaf sampling was conducted randomly while performing PIM. The closest live-leaf to each pin and belonging to one of the five focal species was collected (2-4 leaves per species per subplot). In open-grazed plots in Ny-Ålesund, where vascular plant biomass was very low (see *Results*), leaves were collected at the plot-level. Leaves from each species and subplot were stored in a separate tea-filter bag, pressed dry between filter papers for 72 h, and then oven-dried at 60 °C for 48 h, following Petit Bon, Böhner, et al., (2020). In Semmeldalen, where we did not have access to oven facilities, leaves were kept as dry as possible by regularly substituting filter papers and oven-dried within 5 days.

Each leaf was analysed for C and N concentration with near-infrared reflectance spectroscopy (NIRS FieldSpec 3; ASD Inc.^®^, Boulder, Colorado, USA) in 350− 2500 nm range and using a 4-mm light-adapter for full-leaf analysis (Petit Bon, Böhner, et al., 2020). For each leaf, 3-6 NIR-measurements were taken, depending on leaf size. Each measurement was converted to C and N concentration using prediction models developed for ground-leaf samples (Ancin-Murguzur et al., 2019) and correcting for measures on full leaves (Petit Bon, Böhner, et al., 2020). We computed the median of the measurements of each leaf and then averaged the medians to obtain mean C and N concentration for each focal species within a subplot (for a similar approach, see Petit Bon, Inga, et al., 2020). This led to a total of 66 and 120 leaf nutrient samples from Ny-Ålesund and Semmeldalen, respectively (Appendix 1, Table S2).

We calculated total live-leaf C and N pools:

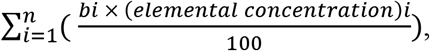

where *n* equals the focal vascular plant species (see above), *b*_*i*_ is the absolute (g m^-2^ dw) live aboveground biomass contributed by species *i*, and *(elemental concentration)*_*i*_ represents either C or N concentration (%dw) of species *i*.

Five-to-ten moss shoots were collected at each interception of the 15 pins (see above) with the moss-mat, and shoots from the same subplot were pooled together. For each moss sample, we separated green and brown parts. Moreover, two samples of organic soil (0-5 cm below the brown part of mosses) were collected from the two corners of each subplot and pooled together. Soil samples were mixed, homogenised and sieved at 2-mm mesh-size.

Moss (green and brown parts separately) and soil samples were analysed with NIRS for C and N concentrations using prediction models developed specifically for mosses (Appendix 1, Figure S1) and soil (Petit Bon, Inga, et al., 2020). We took 3-8 NIR-measurements for each moss sample and 6 NIR-measurements for each soil sample. After converting each measurement to C and N concentration, we computed the average C and N concentrations for mosses and organic soil in each subplot (Appendix 1, Table S2).

#### 2.3.4 Instantaneous ecosystem CO_2_-fluxes

A set of CO_2_-flux measurements was taken twice in each subplot, resulting in a total of 135 and 80 sets from Ny-Ålesund and Semmeldalen, respectively. Each set consisted of a light and a dark measurement, from which we calculated net ecosystem exchange (NEE) and ecosystem respiration (ER), respectively. In the short-term exclosures in Ny-Ålesund, only ER was measured. Estimates of gross ecosystem photosynthesis (GEP) were obtained by subtracting ER from NEE. The two sets of measurements at each subplot were taken on separate days to capture a wider range of environmental (temperature and light) conditions (see below). However, to minimise differences in environmental conditions among measurements taken across treatments (see below), each pair of exclosure and open-grazed plots were measured concurrently.

CO_2_-flux measurements were taken using a closed static/non-steady state system composed of a clear chamber (25 cm × 25 cm area × 35 cm height, made of LEXAN^©^ polycarbonate; >90% light-transmittance), connected to a CO_2_ infrared gas analyser (LI-840A; LICOR Inc.^®^, Lincoln, Nebraska, USA) through an air pump with 0.9-1.0 l min^-1^ flow rate (L052C-11; Parker Corp.^®^, Cleveland, Ohio, USA). To prevent disturbance to the tundra, mattress-foam fabric (1.5 cm thick) was attached to the bottom of the chamber. To minimize air exchange with the external environment, a plastic skirt (30 cm wide) was attached to the bottom of the chamber and held down (i.e. sealing) during measurements by a 2-m long draped around steel chain weighing 4 kg. A fan mixed the air inside the chamber continuously.

Measurements at each subplot took place within 3 h either side of solar noon, started 15-20 s after sealing (acclimation period), and lasted 120-150 s. Air CO_2_ concentration inside the chamber was registered every 5 s. During ER measurements, a completely dark hood was placed over the chamber to exclude light and temporarily halt photosynthesis. At each plot, we first took NEE measures in all subplots, followed by ER measures. Photosynthetically active radiation (PAR) and air temperature (ca. 20 cm above the vegetation) were recorded simultaneously with the CO_2_-fluxes within the chamber every 5 and 10 s, respectively, using a PAR sensor attached to a datalogger (LI-190SA and LI-1400; LICOR Inc.) and a temperature logger (DS1922L-F5; Maxim Integrated^®^, San Jose, California, USA). CO_2_ data were collected under a wide range of light and temperature conditions. PAR during NEE measures was on average 525 ± 256 SD µmol m^-2^ s^-1^ (range: 129-1131 µmol m^-2^ s^-1^), whereas air temperature during both NEE and ER measures was on average 11.4 ± 5.0 °C (5.4-22.1 °C); no significant differences in PAR and air temperature were detected between treatment measures (Appendix 1, Table S3).

Because small-scale spatial variations in hydrological conditions affect ecosystem CO_2_-fluxes in the high-Arctic (Sjögersten et al., 2006), soil volumetric water content (i.e. soil moisture %volume; 0-5 cm depth) was measured in two corners of each subplot after each set of measurements using a hand-held moisture logger (ML3 Theta Probe and HH2 Meter; Delta-T Ltd.^®^, Cambridge, UK). We concurrently measured soil temperature (5 cm depth) in the same two corners using a temperature probe (TFX 410-1 Handheld Thermometer; Ebro^®^, Hagen, Germany).

We used linear regression models to determine ecosystem CO_2_-fluxes (NEE and ER) at each subplot (for a similar approach, see Sundqvist et al., 2020):

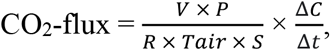

where *V* is the volume of the chamber (0.021875 m^3^), *P* is the average air pressure (atm, registered at Adventdalen weather station [∼10 m a.s.l.] every 1 s and used for both sites), *R* is the ideal gas constant (8.314 J mol^-1^ K^-1^), *T*_*air*_ is the average air temperature (K), *S* is the surface area of the chamber (0.0625 m^2^), and Δ*C/*Δ*t* is the slope of the linear regression, that is, the change in CO_2_ concentration (Δ*C*) within the chamber during the measurement period (Δ*t*). Fluxes are shown as µmol CO_2_ m^-2^ s^-1^ and presented from an atmospheric perspective, namely positive and negative values signify CO_2_ release (C source) and CO_2_ uptake (C sink), respectively, by the ecosystem.

#### 2.4 Statistical analyses

The two experiments were analysed separately. In Ny-Ålesund, there were three levels of ‘herbivory’ (open-grazed [Open], 4-yr-old exclosures [Short-excl], and 15-yr-old exclosures [Long-excl]), whilst in Semmeldalen there were two levels of ‘herbivory’ (open-grazed [Open] and 21-yr-old exclosures [Long-excl]). Pairs of exclosure and open-grazed plots were considered the inferential units (i.e. true replicates; N=5 in both experiments), while subplots within plots were considered the experimental units (per treatment: Ny-Ålesund, n=15; Semmeldalen, n=20). Therefore, if not otherwise described, whenever multiple measures for a given variable were taken in a subplot (e.g. moss-layer depth recorded in the two corners) data were averaged prior to analyses.

We employed a variety of statistical models within a linear mixed-effects model (LMM) framework to account for the spatial structure of the study design, whereby we consistently specified ‘plot’ nested in ‘pair’ as random-effects. ‘Herbivory’ was used as a fixed-effect in all models. Data were *log*-or *arcsine*-transformed when necessary to improve normality and homoscedasticity in model residuals.

#### Vegetation

Changes in vascular plant community structure were assessed on both absolute and relative PFT biomass. For relative changes, we determined proportional biomass by summing the biomass of all PFTs and then calculating the relative contribution of each PFT. Thus, absolute changes are indicative of actual biomass changes, whilst relative changes are indicative of changes in PFT relative abundance. First, we assessed the effect of ‘herbivory’ on absolute total biomass and relative PFT biomass using LMMs and permutational multivariate analysis of variance, respectively. Then, to test for the effects of ‘herbivory’ on absolute biomass of each PFT, we used LMMs fitted separately for each PFT.

LMMs were also built to assess the effects of ‘herbivory’ on standing-dead graminoid biomass and total moss-layer depth. Changes in moss-layer depth were also evaluated separately for the green and brown parts.

#### Ecosystem C and N contents

To investigate how herbivory affects plant and soil chemistry, we used LMMs with either N concentration or C:N as response variables. We fitted separate models for the five focal vascular plant species belonging to the five broadly-classified PFTs, green and brown parts of mosses, and organic soil. Similarly, LMMs were built to assess the effects of herbivory on live-leaf C and N pools.

#### Ecosystem CO_2_-fluxes

To examine how herbivory affects ecosystem CO_2_-fluxes, we used separate LMMs with NEE, ER, or GEP as response variables. Abiotic covariates known to influence CO_2_-fluxes were also incorporated to improve model fit and allow more meaningful ecological interpretations of herbivore effects. We initially included PAR and soil moisture for models of NEE and GEP and soil moisture and soil temperature for models of ER (cf. Sjögersten et al., 2008; Leffler et al., 2019). As in Ny-Ålesund the open-grazed tundra was moss-dominated (see *Results*), and mosses have lower light requirements than vascular plants (Douma et al., 2007), we included a ‘herbivory × PAR’ interaction in NEE and GEP models built on data from there. As well, soil moisture was high and manifested little variation across plots (88% ± 15%; Table S3). Hence, we did not include soil moisture in models from Ny-Ålesund. Finally, as soil moisture influences soil temperature, and these were strongly correlated in Semmeldalen (Appendix 1, Figure S2), we excluded here soil temperature from the ER model to avoid spurious results. Only abiotic covariates that explained a significant proportion of the variance were included in the final models reported in the *Results*. As two sets of CO_2_-flux measurements were taken at each subplot, we here specified ‘subplot’ as an additional nested random-effect in all models to account for the repeated measures.

To gain a better mechanistic understanding of how herbivores affect ecosystem CO_2_-fluxes, we explored the relationships between NEE, ER, or GEP (used in separate LMMs as response variables) and either live vascular plant biomass or moss-layer depth (used as predictors). In these models, we did not include ‘herbivory’ as a fixed-effect as it would be highly correlated with both these predictors. The random structure of the models matched that specified above.

Statistically significant differences between herbivory treatments were defined as having their 95% confidence intervals (CI) not crossing zero. Analogously, significant relationships were also defined as having the 95% CI of their slope not crossing zero. Model summaries are presented in Appendix 1, Tables S4-S11. We performed all data analyses in R ver. 4.2.1 (https://www.r-project.org) using the packages *vegan* (Oksanen et al., 2020), *nlme* (Pinheiro et al., 2015), and *emmeans* (Lenth 2021).

## 3 RESULTS

### 3.1 Vegetation: vascular plants and mosses

In Ny-Ålesund, the experimental exclusion of herbivores (i.e. geese) for 4 and 15 years resulted in 16-fold and 39-fold greater, respectively, live aboveground vascular plant biomass compared to open-grazed tundra (Figure 2a). While either exclosure type had higher biomass of grasses and horsetails compared to open-grazed tundra, only short-term exclosures had higher forb biomass. Live biomass was higher (2×) in long-term than short-term exclosures, and this was due to an 8-fold increase in horsetail biomass. Herbivory treatments also differed in plant-community composition, that is, plant functional type (PFT) relative abundances (Figure 2b). Forbs, grasses and horsetails contributed overall equally to the little vascular plant biomass found in open-grazed tundra. Conversely, short-term and long-term exclosures were in two different vegetation states, dominated by grasses and horsetails, respectively. Goose exclosures also had 120-200-fold higher dead graminoid biomass than open-grazed tundra (0.12 ± 0.52 SE g m^2^), but no differences were detected between short-term (22 ± 11 g m^2^) and long-term (31 ± 15 g m^2^) exclosures (Appendix 1, Table S4).

**FIGURE 2.**
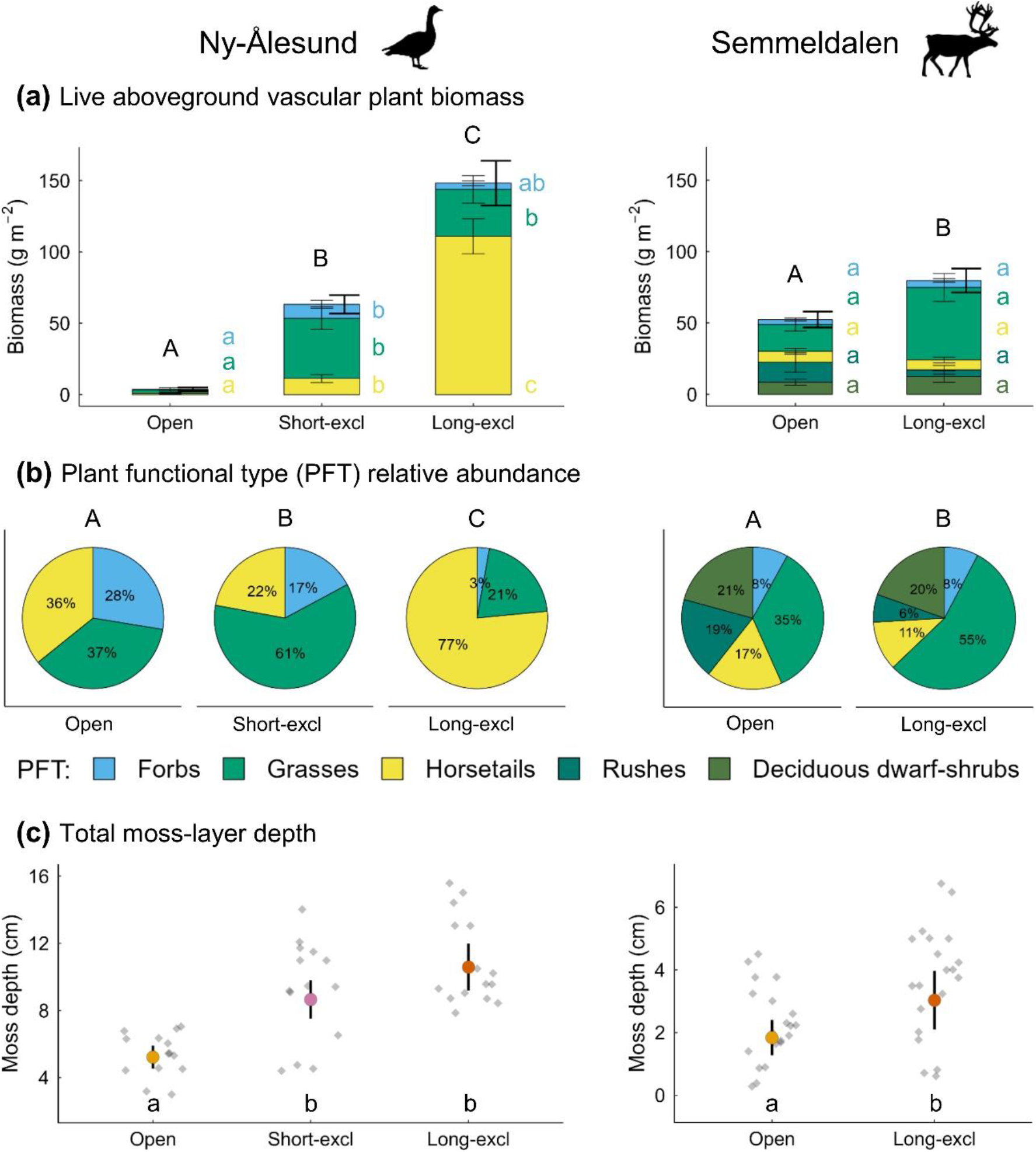
Effects of herbivore removal on vegetation in Ny-Ålesund (left panels) and Semmeldalen (right panels). **(a)** Live aboveground vascular plant biomass in each herbivory treatment (Open: open-grazed; Short-excl: short-term exclosures; Long-excl: long-term exclosures), sorted according to PFTs. Thick and thin bars are the standard errors (SEs) of the mean for total biomass and for the biomass of each PFT, respectively. **(b)** Aboveground PFT relative abundance within each herbivory treatment. **(c)** Model predictions ± SE for total moss-layer depth within each herbivory treatment. Grey dots show raw values; note the different scale of *y*-axis. Treatments connected by different letters indicate a statistically significant difference.

In Semmeldalen, the experimental exclusion of herbivores (i.e. reindeer) for 21 years resulted in 45% greater live aboveground vascular plant biomass compared to open-grazed tundra (Figure 2a). However, the biomass of each individual PFT did not differ between herbivory treatments. Yet, the two treatments still differed in plant-community composition due to a higher relative abundance of grasses and a lower relative abundance of rushes in exclosures compared to open-grazed tundra (Figure 2b). This difference would go unnoticed if these PFTs had been combined as graminoids (Appendix 1, Table S5). Reindeer exclosures also had higher biomass (3×) of dead graminoids (31 ± 17 g m^2^) compared to open-grazed tundra (10 ± 6 g m^2^) (Appendix 1, Table S4).

In Ny-Ålesund, the total moss-layer was almost twice as deep in exclosures than in open-grazed tundra (Figure 2c). This difference was due to contributions from photosynthetically-active green and nearly-decomposed brown parts of the moss-layer, both of which were deeper in short-term (green: 2.1 ± 0.3 cm; brown: 6.3 ± 1.1 cm) and long-term (green: 1.8 ± 0.3 cm; brown: 8.6 ± 1.5 cm) exclosures than in open-grazed tundra (green: 1.0 ± 0.2 cm; brown: 4.0 ± 0.7 cm) (Appendix 1, Table S6).

In Semmeldalen, exclosures had about 60% deeper total moss-layer than open-grazed tundra (Figure 2c). This was due to the depth of the green part being greater within exclosures (1.2 ± 0.3 cm *vs* 0.6 ± 0.1 cm), as that of the brown part did not differ between treatments (Appendix 1, Table S6).

### 3.2 Ecosystem C and N contents

Overall, N concentration and C:N of the three ecosystem compartments (vascular plant leaves, green and brown parts of mosses, and organic soil) displayed relatively few responses to the exclusion of herbivores, and these were only detected in Ny-Ålesund. The grass *Poa arctica* growing in short-term exclosures had about 20% lower N concentration and 30% higher C:N than *Poa arctica* found in open-grazed tundra or in long-term exclosures (Figure 3a,b). Similarly, the green part of mosses growing in short-term exclosures had about 13% lower N concentration (1.19% ± 0.09%) and 17% higher C:N (38.5 ± 3.1) compared to open-grazed tundra (N: 1.38% ± 0.11%; C:N: 31.8 ± 2.5), and tended to have lower N concentration and higher C:N than in long-term exclosures (N: 1.38% ± 0.12%; C:N: 33.2 ± 2.9). Soil N concentration in long-term exclosures (0.87% ± 0.03%) also tended to be lower than in open-grazed tundra (0.96% ± 0.03%) (Appendix 1, Table S8).

**FIGURE 3.**
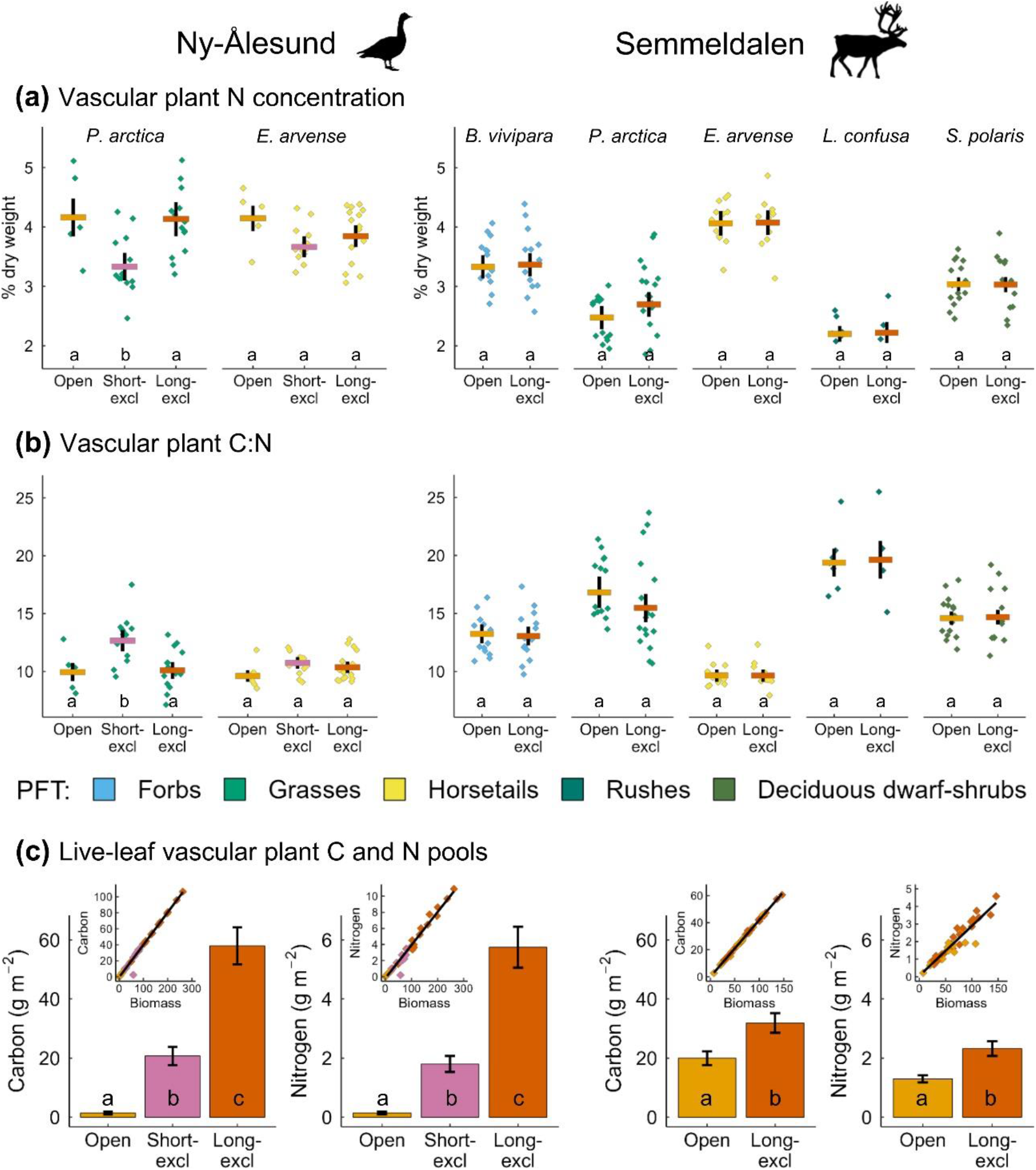
Effects of herbivore removal on vascular plant chemistry in Ny-Ålesund (left panels) and Semmeldalen (right panels). Model predictions ± SE for leaf **(a)** N concentration and **(b)** C:N in each herbivory treatment (Open: open-grazed; Short-excl: short-term exclosures; Long-excl: long-term exclosures) for the five focal vascular plant species. Dots show raw values, coloured according to PFTs. **(c)** Live-leaf vascular plant C and N pools in each herbivory treatment. Bars are the SE of the mean. Insets show the correlation between live aboveground vascular plant biomass and leaf C and N pools. Treatments connected by different letters indicate a statistically significant difference.

Overall, live-leaf C and N pool responses to herbivore removal were similar in their relative magnitude to vascular plant biomass responses, reflecting their tight positive correlations (Figure 3c). In Ny-Ålesund, the exclusion of geese for 4 and 15 years resulted in 13-fold and 40-fold greater, respectively, leaf C and N pools compared to open-grazed tundra. C and N pools also differed between exclosures, with long-term exclosures holding higher (3×) leaf C and N pools than short-term exclosures. In Semmeldalen, reindeer exclusion for 21 years resulted in 60% and 80% greater leaf C and N pools, respectively, compared to open-grazed tundra.

### 3.3 Instantaneous ecosystem CO_2_-fluxes

In Ny-Ålesund, the exclusion of geese for 15 years resulted in 3-fold greater gross ecosystem photosynthesis (GEP), that is, more negative, and ecosystem respiration (ER), that is, more positive, compared to open-grazed tundra (Figure 4a,b). Yet, because of higher GEP than ER, the net CO_2_ exchange of the ecosystem (NEE) was also greater (4×), that is, more negative, in long-term exclosures than in open-grazed tundra (Figure 4c). ER in short-term exclosures was 90% higher and 30% lower than in open-grazed tundra and long-term exclosures, respectively (Figure 4b).

**FIGURE 4.**
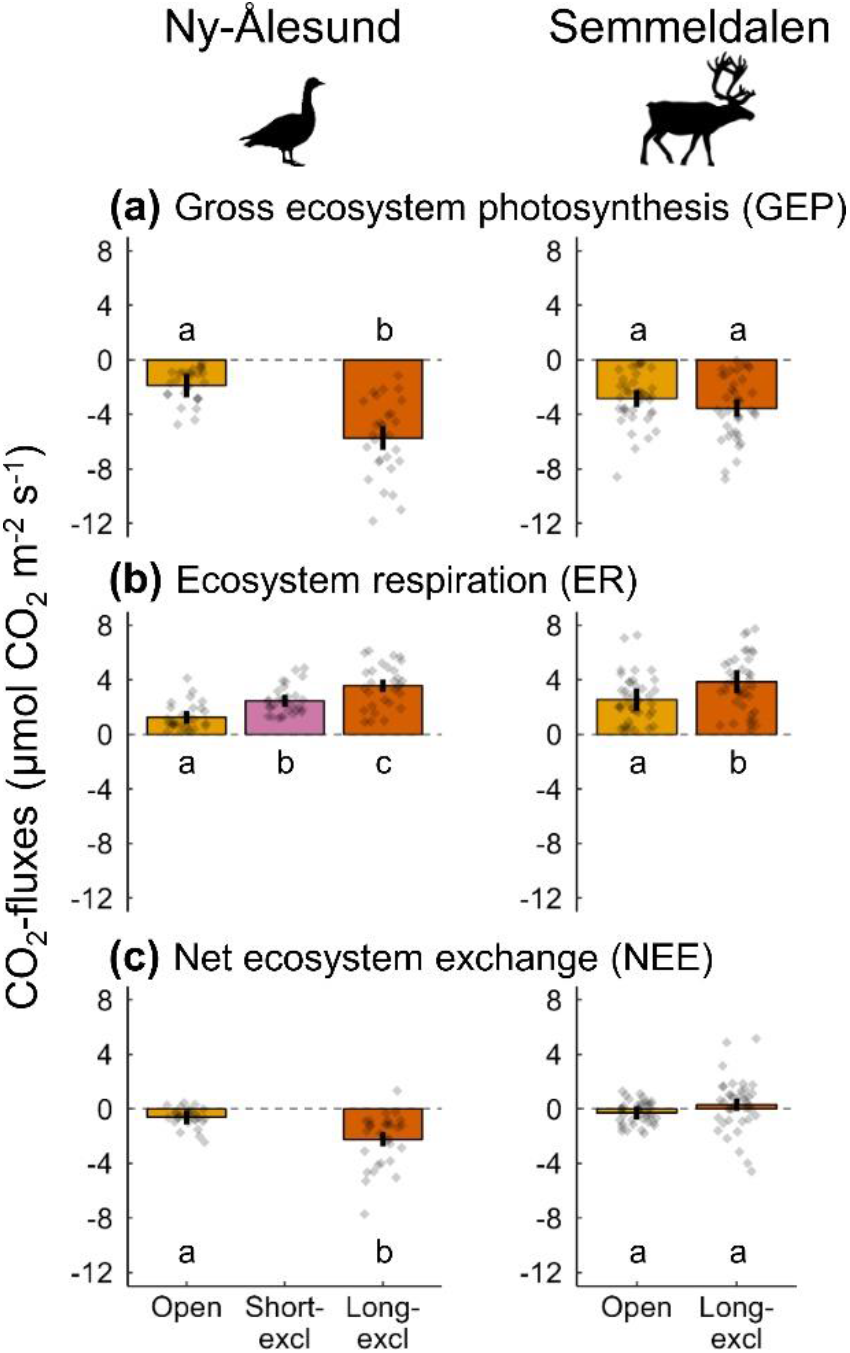
Effects of herbivore removal on ecosystem CO_2_-fluxes in Ny-Ålesund (left panels) and Semmeldalen (right panels). Model predictions ± SE for **(a)** GEP, **(b)** ER, and **(c)** NEE in each herbivory treatment (Open: open-grazed; Short-excl: short-term exclosures; Long-excl: long-term exclosures); final models included the abiotic covariates that explained a significant proportion of the variance (Figure 5a; see *Material and Methods: Statistical analyses*). Dots show raw values. Treatments connected by different letters indicate a statistically significant difference.

In Semmeldalen, the exclusion of reindeer for 21 years increased ER by 50% compared to open-grazed tundra (Figure 4b), but neither affected GEP nor NEE (Figure 4a,c). Yet, it is noteworthy that, although no differences in NEE were detected, exclosures showed a weak net loss of CO_2_, whilst open-grazed tundra showed a weak net uptake of CO_2_ (Figure 4c).

Overall, photosynthetically active radiation (PAR) was a significant abiotic predictor of the variation in GEP and NEE in Ny-Ålesund and GEP in Semmeldalen (Figure 5a). However, in Ny-Ålesund, the explanatory power of PAR depended on herbivory treatment, indicating that both flux components were related to PAR in long-term exclosures, but not in open-grazed tundra. In Semmeldalen, soil moisture was a good predictor of NEE, with the ecosystem switching from net loss to net uptake of CO_2_ above about 50% soil moisture content (Figure 5a).

**FIGURE 5.**
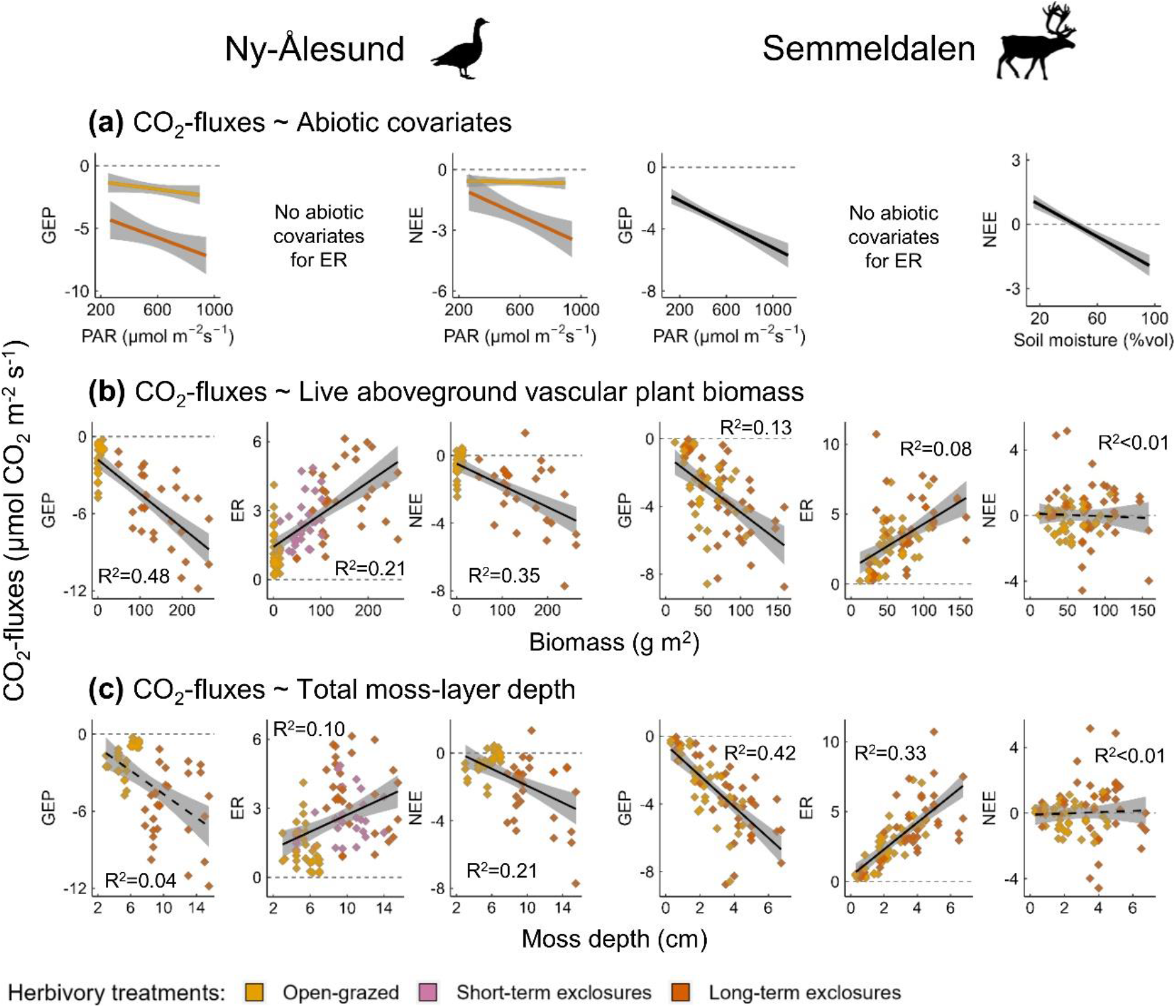
Relationships between ecosystem CO_2_-fluxes and environmental variables in Ny-Ålesund (left panels) and Semmeldalen (right panels). **(a)** CO_2_-fluxes (GEP, ER, and NEE) in relation to changes in abiotic covariates that explained a significant proportion of the variance and were thus included in the final models of herbivore removal effects on CO_2_-fluxes (Figure 4; see *Material and Methods: Statistical analyses*). Relationships between GEP or NEE and PAR in Ny-Ålesund are displayed separately for each herbivory treatment (coloured lines), as the ‘herbivory × PAR’ interaction was retained in the final model. CO_2_-fluxes as predicted by **(b)** live aboveground vascular plant biomass and **(c)** total moss-layer depth (dots show raw values, coloured according to treatments). Statistically significant and non-significant relationships are displayed with solid and dashed lines, respectively (marginal R^2^ are shown in each panel). Lines and bands represent regression lines and their 95% CIs. Note the different scale on the *y-* and *x-*axis depending on which CO_2_-flux component and predictor, respectively, is displayed.

Overall, GEP, ER, and NEE were linked to both live aboveground vascular plant biomass (Figure 5b) and total moss-layer depth (Figure 5c) at both the Ny-Ålesund and Semmeldalen sites. Exceptions were the relationship between GEP and moss-layer depth in Ny-Ålesund and relationships with NEE in Semmeldalen, which were weak.

Soil temperature decreased with increasing total moss-layer depth in Semmeldalen, but not in Ny-Ålesund, whereas soil moisture was not related to moss depth at either site (Appendix 1, Figure S3).

## 4 DISCUSSION

In this study, we aimed at quantifying how high-Arctic ecosystems respond to the long-term removal of migratory geese and resident, non-migratory ungulates, two dominating tundra herbivores. Standardized data from a wet habitat (wetland vegetation) utilised by barnacle geese (Ny-Ålesund site) and from mesic-to-dry habitats (meadow/heath vegetation) utilised by reindeer (Semmeldalen site) revealed that both herbivores influence tundra vegetation and ecosystem CO_2_-fluxes. Whilst the effects of geese were larger than those exerted by reindeer, this difference is scale-dependent, with geese being more spatially concentrated and affecting a smaller portion of the tundra landscape compared to the more widely dispersed reindeer.

Our findings indicate that functionally different herbivores with contrasting spatial, temporal and feeding ecology differ in the extent they affect their main habitat utilised for foraging. This highlights the substantial heterogeneity in long-term herbivore impacts on ecosystem functioning, with important implications in relation to rapidly changing herbivore populations in a warming Arctic.

In the wet habitat in Ny-Ålesund, the presence of migratory geese and their exclusion for 4 and 15 years led to three different vegetation states: moss-dominated (heavily grazed), graminoid-dominated, and horsetail-dominated tundra, respectively (cf. Ravolainen et al., 2020). This shows that, when concentrated in space (in our case due to them having flightless goslings and/or wing feather moult), and even for a relatively short period of time around peak summer, geese can influence tundra wetlands by maintaining low levels of standing aboveground vascular plant biomass (Sjögersten et al., 2008; Kuijper et al., 2009), and in turn considerably reduce vegetation C and N pools (our study; Sjögersten et al., 2011). Moreover, goose-grazed tundra had almost no standing-dead graminoid biomass and a moss-layer that was about half the depth of that within exclosures, suggesting that both grazing on mosses (Soininen et al., 2010) and trampling effects (Tuomi et al., 2021) may be considerable. Yet, this wetland appeared to successfully recover following goose exclusion. By using the same experimental set-up as we did, Sjögersten et al., (2011) found that removing geese for 4 years greatly increased grass biomass and its C pool. We extend those results by showing that the system’s potential for recovery is not depleted despite 11 years of additional grazing (results from the newly established 4-year-old exclosures), and that currently the successional trajectory of the vegetation in the longer-term is characterized by a replacement of grasses by horsetails (results from the 15-year-old exclosures); for an overview of the notable increase in horsetail abundance in this part of Svalbard, see Van der Wal et al., (2020). Therefore, our findings indicate that near-irreversible, degraded vegetation states, as those established following grazing and grubbing by expanding populations of lesser snow goose in the Canadian low-Arctic (Jefferies et al., 2006), are not yet occurring in Svalbard, though geese exerting control over its wetlands (Van der Wal & Hessen 2009).

In mesic-to-dry habitats in Semmeldalen, 21 years of reindeer removal resulted in 45% higher standing aboveground vascular plant biomass. This difference between open-grazed tundra and exclosures contrasts with the greater effects of geese on aboveground biomass in Ny-Ålesund. Although Semmeldalen and the adjacent valley system host one of the densest populations of reindeer in Svalbard (6-8 animals per km2; Le Moullec et al., 2019), estimates of grazing pressure on vascular plants indicate overall low offtake rates (Van der Wal et al., 2000; Van der Wal et al., 2004). Low offtake rates have also been observed in another high-Arctic system, in northeast Greenland, with muskoxen, despite at high densities, removing a very low amount of total plant biomass (Mosbacher et al., 2016). This suggests that, although non-migratory, large herbivores utilise the high-Arctic tundra extensively, their impact on vascular plant biomass is typically low, either because distributed on a large spatial scale or because plants are able to at least partly compensate the losses suffered from grazing (but see Mosbacher et al., 2018 for strong vegetation responses to 5-year muskox exclusion in a high-Arctic fen). Importantly, reindeer reduced moss-layer depth, raising soil temperatures in turn (cf. Gornall et al., 2009), and did not affect the relative abundance of graminoids, suggesting the potential for a positive feedback between herbivores and their forage plants despite the resultant decreased biomass (Van der Wal & Brooker 2004; Petit Bon, Inga, et al., 2020). As well, lower standing biomass in open-grazed tundra does not equate to that the ecosystem is found in a less productive state, as a fraction of vegetation, and hence of ecosystem C and N pools, accumulated in herbivore body mass (Bråthen et al., 2021).

We found differences in vegetation (vascular plants and mosses) and soil chemistry between open-grazed tundra and exclosures in the wet habitat in Ny-Ålesund, but not in mesic-to-dry habitats in Semmeldalen. In Ny-Ålesund, N concentration of *Poa arctica*, which accounted for 40% of the aboveground biomass across herbivory treatments, was the lowest (and C:N the highest) in the short-term exclosures, whereas no differences were detected between goose-grazed tundra and long-term exclosures. The same pattern was found for the green part of mosses, indicating that nutrient levels in the vegetation (and the quality of goose forage species) are decreased by short-term goose removal (Gauthier et al., 1995; Beard, Choi, et al., 2019). Two different pathways are likely responsible for the high N concentration (and lower C:N) in the vegetation found in open-grazed tundra and long-term exclosures. The quick return of nutrients in the form of animal excreta and the maintenance of leaves in early phenological stages through repeated grazing (Bazely & Jefferies 1985; Petit Bon, Inga, et al., 2020) might be the short-term mechanisms acting in the presence of geese (i.e. the fast herbivore-mediated pathway). Conversely, in the protracted absence of geese, it is the build-up of a large N pool in vascular plants that eventually regulates the return rate of N to the system (i.e. the slow vegetation-mediated pathway), as vascular plant litter releases nutrients more promptly than moss litter (Fivez 2014). Because the moss-layer was much deeper in exclosures and mosses efficiently absorb nutrients released from litter (Gornall et al., 2009), it is not surprising that soil N tended to be higher in open-grazed tundra compared to long-term exclosures. Overall, these observations point to that whole system’s nutrient dynamics are generally slower in the short-term absence of geese.

The result that 15 years of goose exclusion in Ny-Ålesund caused a 4-fold increase in peak summer net CO_2_ uptake rate of the ecosystem (NEE) aligns with findings from other studies showing that intense herbivory by either geese (Van der Wal et al., 2007; Sjögersten et al., 2011; Leffler et al., 2019) or mammals (Metcalfe & Olofsson 2015) can substantially reduce the C sink strength of tundra ecosystems. Larger gross ecosystem photosynthesis (GEP) than ecosystem respiration (ER), combined with their strong relationships with vascular plant biomass, and to a lesser extent moss-layer depth, indicates that the goose-induced changes in the instantaneous C exchange rates of this high-Arctic wetland were predominantly driven by changes in vegetation as opposed to soil (Sjögersten et al., 2008). This is lent further support by PAR controlling both GEP and NEE, at least at appreciable levels of plant biomass (i.e. within exclosures), whereas soil temperature did not influence ER, likely because microbial respiration was constrained by elevated soil moisture content (Sjögersten et al., 2006). The fact that ER in short-term exclosures was 90% higher than ER in open-grazed tundra, but only 30% lower than ER in long-term exclosures, indicates that the system’s C exchange rates might be characterized by an initially rapid recovery from grazing, after which the recovery rate slows down. This occurred despite strong differences in vegetation composition (as discussed above), suggesting that short-term *vs* long-term goose removal can restructure Arctic wetland plant communities without causing additional major alterations in ecosystem C dynamics (cf. Sistla et al., 2013).

Interesting insights also emerge when comparing our findings to those obtained by Sjögersten et al., (2008), who used captive barnacle geese to achieve short-term experimental herbivory in previously naturally ungrazed habitats in Adventdalen, a valley in central Svalbard. First, CO_2_-fluxes in our exclosures were 2-to-6 times greater than those observed in the ungrazed wet habitat in Adventdalen, which highlight how high-Arctic wetlands may considerably vary in ecosystem process rates. Second, the fact that experimentally controlled goose grazing in previously ungrazed tundra strongly reduced the C sink strength of the wetland (also found in our study), but not of the mesic heath, suggests that the impacts of geese depend upon habitat type, reflecting that they naturally graze more intensively in wet habitats (Speed et al., 2009; Madsen et al., 2011).

Long-term reindeer removal in Semmeldalen caused little change in ecosystem CO_2_-fluxes. Yet, ER was higher in the exclosures than in open-grazed tundra. Because GEP did not increase concurrently, it is likely that the exclosure-induced change in ER was predominantly soil driven. By decreasing plant biomass and thereby plant C pool, long-term reindeer herbivory may have limited the C available for soil microbes, hence decreasing microbial C biomass and microbial respiration (Stark & Grellmann 2002). We found both GEP and ER were tightly related to moss-layer depth, and to a lesser extent vascular plant biomass, which highlights the dominant, direct contribution of mosses to instantaneous C exchange rates of mesic-to-dry high-Arctic habitats (cf. Douma et al., 2007). Peculiarly, moss-GEP and moss-ER relationships were stronger here than in the wet habitat in Ny-Ålesund, despite its deeper moss-layer, suggesting there might be a depth-threshold after which mosses exert their maximum control on GEP and ER. Results from Semmeldalen also reveal that, because GEP and ER concurrently increased at a similar rate with increasing moss-layer depth, and hence cancelled each other out, mosses did not exert a detectable influence on NEE. Instead, we found NEE to be controlled by soil moisture (cf. Sjögersten et al., 2006), which did not relate to moss-layer depth. Combined, these observations lead to two important considerations for the C dynamics of high-Arctic meadows/heaths utilised by large herbivores. If herbivore populations continue to increase (Schmidt et al., 2015; Le Moullec et al., 2019; Solberg et al., 2022), further lowering moss-layer depth and vascular plant biomass, alterations in GEP and ER should be expected, though the NEE may remain relatively uninfluenced. However, should such an additional decrease in moss-layer depth translate into enhanced soil evaporation rates (Blok et al., 2011), grazed mesic-to-dry habitats could eventually turn into a C source. This might be exacerbated if the decreased moss-layer depth or lower soil moisture boost soil temperatures (our study; Gornall et al., 2009; Blok et al., 2011), eventually increasing ER by stimulating soil respiration, as long as the resulting lower C pool does not become limiting for soil microbes.

## 5 CONCLUSIONS

The present study emphasises the heterogeneity of tundra ecosystem responses to herbivory. Our evaluation of long-term effects of herbivore removal on vegetation, ecosystem C and N contents and instantaneous CO_2_-fluxes, albeit limited to the peak of the growing season, shows that geese, highly aggregated in space and time, exert a stronger effect on Svalbard’s high-Arctic tundra compared to more widely dispersed reindeer. The different effect of these two herbivores seen in their preferred habitat reflects natural differences in habitat sensitivity and habitat utilisation, as well as the fact that herbivory varies spatially due to the heterogeneous distribution of forage and other constraints, such as the availability of safe grounds where geese undergo full wing moult and goslings grow up, which limit their spatial movement. Therefore, we show that some habitats can be strongly impacted by herbivores, whereas in other habitats their influence is likely to be minor.

Due to the lack of parallel, similarly designed, long-term herbivore removal experiments in the very same system, studies such as ours will remain an exception. Yet, we believe that assessing how functionally different herbivores affect ecosystems, in light of the spatial and temporal dimensions of the way they use the landscape, will help us understand the role of herbivory for processes and functions of the habitats most extensively utilised for foraging. This, in turn, will provide vital information to project the effects of changing herbivore population densities on ecosystem functioning, and refine our predictions on whether these changes are likely to mitigate or further amplify the impact of climate change.

## Data availability

Data will be available at the NSF Arctic Data Center Repository, upon acceptance for publication of this Manuscript.

## Authors’ contributions

MPB, BBH, RvdW, and MJJEL had the initial idea, which was discussed with all Authors at different stages in the order: ISJ, MLM, AP, KAB, KLM, HB, and KHB. RvdW and MJJEL originally designed the long-term experiments. MPB, RvdW, MJJEL, and AP planned methodology. MPB, AP, and KLM collected the data. HB and MPB wrote the R-script for CO_2_ data processing. KAB and HB developed NIRS models for mosses. MPB processed the samples, analysed the data, and wrote the manuscript, to which RvdW and BBH provided key contributions. All Authors commented on previous drafts and approved the final version for publication. There are no conflicts of interest to declare.

## Acknowledgments

This research was financed by the Research Council of Norway (FRIPRO grant 276080, to BBH). Additional funding for field material was equally provided by The Governor of Svalbard (Svalbard Environmental Protection Fund, grant 15/128) and the Research Council of Norway (Arctic Field Grant, grant 269957) (both to MPB). MPB is currently supported by the National Science Foundation (grant ANS-2113641, to KHB). The Governor of Svalbard gave permit for fieldwork and sampling (RiS-IDs 6359 and 3577). We thank the Logistic Department at UNIS and staff at COAT (Climate-Ecological Observatory of the Arctic Tundra) for providing technical support, and UiT for providing Lab facilities. We are grateful to Benjamin Segger for field assistance in Ny-Ålesund. Finally, we thank the numerous field assistants and technicians that, throughout the years, helped in establishing and maintaining the long-term experiments.

