## Supporting Information for "Long-term herbivore removal experiments reveal different impacts of geese and reindeer on vegetation and ecosystem CO_2_-fluxes in high-Arctic tundra"

Below are the supplementary tables, figures, and results supporting the study "**Long-term herbivore removal experiments reveal different impacts of geese and reindeer on vegetation and ecosystem $CO_2$-fluxes in high-Arctic tundra**".

Matteo Petit Bon[1,2*], Brage B. Hansen[3,4], Maarten J. J. E. Loonen[5], Alessandro Petraglia[6], Kari Anne Bråthen[7], Hanna Böhner[2,7], Kate Layton-Matthews[4,8], Karen H. Beard[1], Mathilde Le Moullec[4], Ingibjörg S. Jónsdóttir[2,9], René van der Wal[10]

[1] Department of Wildland Resources | Quinney College of Natural Resources and Ecology Center, Utah State University, UT-84322 Logan, Utah, USA

[2] Department of Arctic Biology, University Centre in Svalbard, N-9171 Longyearbyen, Norway

[3] Norwegian Institute for Nature Research, NO-7034 Trondheim, Norway

[4] Centre for Biodiversity Dynamics | Department of Biology, Norwegian University of Science and Technology, NO-7491 Trondheim, Norway

[5] Arctic Centre, University of Groningen, CW-9718 Groningen, Netherlands

[6] Department of Chemistry, Life Sciences, and Environmental Sustainability, University of Parma, IT-43124 Parma, Italy

[7] Department of Arctic and Marine Biology | Faculty of Biosciences, Fisheries, and Economics, Arctic University of Norway, N-9037 Tromsø, Norway

[8] Norwegian Institute for Nature Research, Fram Centre, N-9296 Tromsø, Norway

[9] Institute of Life and Environmental Sciences, University of Iceland, IS-102 Reykjavik, Iceland

[10] Department of Ecology, Swedish University of Agricultural Sciences, SE-75651 Uppsala, Sweden

ORCID-ID: https://orcid.org/0000-0001-9829-8324

BBH: https://orcid.org/0000-0001-8763-4361

MJJEL: https://orcid.org/0000-0002-3426-4595

AP: https://orcid.org/0000-0003-4632-2251

HB: https://orcid.org/0000-0001-7356-5457

KAB: https://orcid.org/0000-0003-0942-1074

KLM: https://orcid.org/0000-0001-5275-1218

KHB: https://orcid.org/0000-0003-4997-2495

MLM: https://orcid.org/0000-0002-3290-7091

ISJ: https://orcid.org/0000-0003-3804-7077

RvdW: https://orcid.org/0000-0002-9175-0266

### APPENDIX 1

#### Supplementary Tables

**TABLE S1.** Vascular plant species and corresponding plant functional types. Mean relative (%) abundance (and standard deviation) of vascular plant species (or genera when species were not further identified) in the long-term **(a)** goose removal experiment (Ny-Ålesund) and **(b)** reindeer removal experiment (Semmeldalen), separated for the three herbivory treatments (open-grazed tundra, short-term exclosures and long-term exclosures); species/genera are listed in alphabetical order within their plant functional types. Note that (b) in Semmeldalen, no short-term exclosures were available (*Material and Methods: Experimental sites and setup*). Species/genus names follow the Svalbard Flora (https://www.svalbardflora.no).

| | Plant Functional Type (PFT) | Plant species | Percent Relative Abundance (%) | | |
|---|---|---|---|---|---|
| | | | Open-grazed tundra | Short-term exclosures | Long-term exclosures |
| **(a)** Ny-Ålesund (goose exclusion) | Forbs | *Cardamine* spp. (*C. bellidifolia* and *C. nymanii*) | 15.3 (26.8) | 11.7 (15.3) | 2.2 (3.1) |
| | | *Ranunculus* spp. (*R. nivalis* and *R. glacialis*) | 3.2 (12.1) | 0 (0) | 0 (0) |
| | | *Saxifraga cernua* | 0 (0) | 5.5 (10.1) | 0.7 (1.8) |
| | | *Saxifraga cespitosa* | 9.1 (14.9) | 0 (0) | 0 (0) |
| | Grasses | *Poa arctica* | 36.6 (36.0) | 60.8 (32.4) | 20.6 (18.8) |
| | Horsetails | *Equisetum arvense* | 35.8 (27.6) | 22.1 (23.5) | 76.5 (18.3) |
| **(b)** Semmeldalen (reindeer exclusion) | Forbs | *Bistorta vivipara* | 6.6 (8.5) | | 6.2 (9.2) |
| | | *Cerastium arcticum* | 0 (0) | | 0.3 (1.3) |
| | | *Cerastium* spp. (*C. cerastoides* and *C. alpinum*) | 0.2 (0.7) | | 1.1 (3.6) |
| | | *Draba* spp. | 0.04 (0.1) | | 0 (0) |
| | | *Koenigia islandica* | 0.6 (2.2) | | 0 (0) |
| | | *Micranthes nivalis* | 0.04 (0.2) | | 0 (0) |
| | | *Oxyria digyna* | 0.2 (1.0) | | 0 (0) |
| | | *Ranunculus hyperboreus* | 0.2 (0.6) | | 0.1 (0.5) |
| | | *Sagina nivalis* | 0.3 (1.1) | | 0 (0) |
| | | *Saxifraga oppositifolia* | 0.03 (0.1) | | 0 (0) |
| | | *Stellaria longipes* | 0 (0) | | 0.01 (0.1) |
| | Grasses | *Alopecurus ovatus* | 4.5 (8.8) | | 2.1 (4.8) |
| | | *Calamagrostis neglecta* | 0 (0) | | 0.2 (0.9) |
| | | *Festuca rubra* | 3.9 (6.5) | | 0.8 (2.1) |
| | | *Poa arctica* | 26.9 (27.8) | | 52.0 (32.9) |
| | Horsetails | *Equisetum arvense* | 17.3 (18.0) | | 11.2 (16.5) |
| | Rushes | *Juncus biglumis* | 0.2 (0.7) | | 0 (0) |
| | | *Luzula confusa* | 18.0 (35.5) | | 6.3 (16.9) |
| | | *Luzula nivalis* | 0.5 (2.0) | | 0 (0) |
| | Deciduous dwarf-shrubs | *Salix polaris* | 20.8 (22.9) | | 19.6 (26.7) |

**TABLE S2.** Overview of the plant and soil samples analysed for carbon and nitrogen concentrations. Number of samples for vascular plants (five focal species), mosses (green and brown parts), and organic soil collected in the long-term **(a)** goose removal experiment (Ny-Ålesund) and **(b)** reindeer removal experiment (Semmeldalen), separated for the three herbivory treatments (open-grazed tundra, short-term exclosures and long-term exclosures). Note that (b) in Semmeldalen, no short-term exclosures were available (*Material and Methods: Experimental sites and setup*).

| | **Sample type** | **Open-grazed tundra** | **Short-term exclosures** | **Long-term exclosures** |
|---|---|---|---|---|
| **(a)** Ny-Ålesund (goose exclusion) | *Poa arctica* (grass) | 5 | 13 | 15 |
| | *Equisetum arvense* (horsetail) | 5 | 13 | 15 |
| | Green mosses | 8 | 8 | 5 |
| | Brown mosses | 14 | 13 | 8 |
| | Organic soil | 10 | 10 | 10 |
| **(b)** Semmeldalen (reindeer exclusion) | *Bistorta vivipara* (forb) | 13 | | 14 |
| | *Poa arctica* (grass) | 14 | | 18 |
| | *Equisetum arvense* (horsetail) | 12 | | 11 |
| | *Luzula confusa* (rush) | 7 | | 4 |
| | *Salix polaris* (deciduous dwarf-shrub) | 15 | | 12 |
| | Green mosses | 17 | | 17 |
| | Brown mosses | 20 | | 20 |
| | Organic soil | 20 | | 20 |

**TABLE S3.** Overview of the environmental conditions during $CO_2$-flux measurements and quality of the $CO_2$-flux measurements. Mean ± standard deviation of environmental conditions during $CO_2$-flux measurements and of the quality ($R^2$) of the $CO_2$-flux measurements in the long-term **(a)** goose removal experiment (Ny-Ålesund) and **(b)** reindeer removal experiment (Semmeldalen), separated for the three herbivory treatments (open-grazed tundra, short-term exclosures and long-term exclosures). Note that (a) in Ny-Ålesund, no measurements of NEE were taken in short-term exclosures (*Material and Methods: Sample collection and processing*), and that (b) in Semmeldalen, no short-term exclosures were available (*Material and Methods: Experimental sites and setup*). Abbreviations: PAR = photosynthetically active radiation; NEE = net ecosystem exchange; ER = ecosystem respiration.

| | Environmental conditions and measurement quality | Open-grazed tundra | Short-term exclosures | Long-term exclosures |
|---|---|---|---|---|
| **(a)** Ny-Ålesund (goose exclusion) | PAR NEE (µmol $m^{-2}$ $s^{-1}$) | 576 ± 209 | | 609 ± 226 |
| | Chamber temperature NEE (°C) | 9.0 ± 2.7 | | 9.5 ± 3.0 |
| | Chamber temperature ER (°C) | 8.9 ± 2.7 | 6.9 ± 0.7 | 9.5 ± 2.9 |
| | Soil moisture (%volume) | 90.1 ± 16.5 | 85.2 ± 11.9 | 88.8 ± 17.9 |
| | Soil temperature (°C) | 2.0 ± 1.0 | 2.8 ± 1.5 | 2.4 ± 1.2 |
| | $R^2$ CO2 fluxes (NEE) | 0.90 ± 0.24 | | 0.98 ± 0.02 |
| | $R^2$ CO2 fluxes (ER) | 0.97 ± 0.04 | 0.99 ± <0.01 | 0.99 ± 0.01 |
| **(b)** Semmeldalen (reindeer exclusion) | PAR NEE (µmol $m^{-2}$ $s^{-1}$) | 469 ± 267 | | 484 ± 282 |
| | Chamber temperature NEE (°C) | 14.6 ± 4.9 | | 14.8 ± 5.1 |
| | Chamber temperature ER (°C) | 14.7 ± 4.9 | | 14.7 ± 5.0 |
| | Soil moisture (%volume) | 43.6 ± 20.7 | | 44.5 ± 23.6 |
| | Soil temperature (°C) | 7.7 ± 2.0 | | 7.2 ± 1.7 |
| | $R^2$ CO2 fluxes (NEE) | 0.82 ± 0.33 | | 0.86 ± 0.30 |
| | $R^2$ CO2 fluxes (ER) | 0.98 ± 0.04 | | 0.99 ± 0.01 |

**TABLE S4.** Results from linear mixed-effects models on the effects of herbivore removal on absolute aboveground vascular plant biomass. Parameter estimates of fixed-effects (Est.) and their 95% confidence interval (CI – lower and upper bounds) for the models on aboveground vascular plant biomass (*log-transformed+1*) in the long-term **(a)** goose removal experiment (Ny-Ålesund) and **(b)** reindeer removal experiment (Semmeldalen). The intercept (Int.) refers to the biomass in the baseline herbivory treatment, whereas Est. refers to the change in biomass in the treatment being contrasted relative to the baseline (e.g. in the contrast 'Short-excl vs Open', Int. refers to open-grazed tundra and Est. refers to the change in short-term exclosures relative to Int.). Estimates in bold indicate that their 95% CI does not include zero (i.e. statistically significant effects), whereas estimates in italic indicate that their 90% CI does not include zero (i.e. biologically interesting, close-to-significant effects). Note that (b) in Semmeldalen, no short-term exclosures were available (*Material and Methods: Experimental sites and setup*). Abbreviations: Open = open-grazed tundra; Short-excl = short-term exclosures; Long-excl = long-term exclosures.

**TABLE S4.**

| | Model* | Contrast | Int. | Est. | low CI | up CI | *P-value* |
|---|---|---|---|---|---|---|---|
| **(a)** Ny-Ålesund (goose exclusion) | Live vascular plant (VP) biomass | Short-excl vs Open | 1.09 | **2.98** | **2.07** | **3.88** | 0.0001 |
| | | Long-excl vs Open | 1.09 | **3.83** | **2.92** | **4.73** | <0.0001 |
| | | Long-excl vs Short-excl | 4.07 | **0.85** | **0.05** | **1.76** | 0.0414 |
| | Forb biomass | Short-excl vs Open | 0.20 | **1.70** | **0.74** | **2.66** | 0.0036 |
| | | Long-excl vs Open | 0.20 | *0.84* | *-0.12* | *1.80* | 0.0772 |
| | | Long-excl vs Short-excl | 1.90 | *-0.85* | *-1.81* | *0.11* | 0.0747 |
| | Grass biomass | Short-excl vs Open | 0.65 | **2.58** | **1.22** | **3.94** | 0.0024 |
| | | Long-excl vs Open | 0.65 | **2.26** | **0.90** | **3.62** | 0.0051 |
| | | Long-excl vs Short-excl | 3.23 | -0.32 | -1.68 | 1.04 | 0.6051 |
| | Horsetail biomass | Short-excl vs Open | 0.48 | **1.53** | **0.40** | **2.66** | 0.0142 |
| | | Long-excl vs Open | 0.48 | **4.15** | **3.02** | **5.28** | <0.0001 |
| | | Long-excl vs Short-excl | 2.00 | **2.62** | **1.49** | **3.75** | 0.0007 |
| | Standing-dead graminoid biomass | Short-excl vs Open | 0.11 | **3.03** | **1.73** | **4.33** | 0.0007 |
| | | Long-excl vs Open | 0.11 | **3.37** | **2.07** | **4.67** | 0.0003 |
| | | Long-excl vs Short-excl | 3.14 | 0.34 | -0.96 | 1.64 | 0.5598 |
| **(b)** Semmeldalen (reindeer exclusion) | Live VP biomass | Long-excl vs Open | 3.86 | **0.41** | **0.03** | **0.79** | 0.0411 |
| | Forb biomass | Long-excl vs Open | 1.17 | 0.13 | -0.62 | 0.88 | 0.6515 |
| | Grass biomass | Long-excl vs Open | 2.39 | 1.02 | -0.40 | 2.44 | 0.1177 |
| | Horsetail biomass | Long-excl vs Open | 1.44 | -0.11 | -0.66 | 0.45 | 0.6175 |
| | Rush biomass | Long-excl vs Open | 1.09 | -0.56 | -1.82 | 0.70 | 0.2852 |
| | Deciduous dwarf-shrub biomass | Long-excl vs Open | 1.61 | -0.09 | -1.83 | 1.64 | 0.8882 |
| | Standing-dead graminoid biomass | Long-excl vs Open | 2.38 | **1.10** | **0.09** | **2.11** | 0.0390 |

(*) The models 'Live vascular plant biomass' refer to total live aboveground vascular plant biomass; the models on plant functional types (PFTs; forbs, grasses, horsetails, rushes, and deciduous dwarf-shrubs) refer to live aboveground PFT biomass; the models 'Standing-dead graminoid biomass' refer to standing-dead biomass of graminoids (grasses and rushes), and do not include the litter layer.

**TABLE S5.** Results from permutational multivariate analysis of variance (PERMANOVA) on the effects of herbivore removal on overall plant-community composition. Model outputs for differences in overall plant-community composition (relative aboveground plant functional type abundances) among herbivory treatments (DF = degrees of freedom; SS = sum of squares; MS = mean of squares; $R^2$ = explained variance; F-model = pseudo-F statistics) in the long-term **(a)** goose removal experiment (Ny-Ålesund) and **(b)** reindeer removal experiment (Semmeldalen). PERMANOVA was run on the Bray-Curtis distance matrix (based on the root square-arcsine transformation of relative plant functional type abundances at each subplot) and consisted of 10000 restricted permutations to account for the hierarchical spatial structure of the study design ('subplots' nested within 'plots', in turn nested within 'pairs'; *Material and Methods: Statistical analyses*). *P-values* in bold indicate that overall plant-community composition significantly differed among the herbivory treatments considered in the model. Note that (b) in Semmeldalen, no short-term exclosures were available (*Material and Methods: Experimental sites and setup*). Abbreviation: PFT = plant functional type.

| | Model* | Model term | DF | SS | $R^2$ | F-model | *P-value* |
|---|---|---|---|---|---|---|---|
| **(a)** Ny-Ålesund (goose exclusion) | All herbivory treatments | Herbivory treatment | 2 | 1.364 | 0.300 | 8.776 | **0.0001** |
| | | Residuals | 41 | 3.187 | 0.700 | | |
| | | Total | 43 | 4.552 | 1.000 | | |
| | Open-tundra and short-term exclosures | Herbivory treatment | 1 | 0.242 | 0.080 | 2.362 | **0.0324** |
| | | Residuals | 27 | 2.766 | 0.920 | | |
| | | Total | 28 | 3.008 | 1.000 | | |
| | Open-tundra and long-term exclosures | Herbivory treatment | 1 | 0.648 | 0.233 | 8.198 | **0.0001** |
| | | Residuals | 27 | 2.135 | 0.767 | | |
| | | Total | 28 | 2.783 | 1.000 | | |
| | Short-term exclosures and long-term exclosures | Herbivory treatment | 1 | 1.141 | 0.436 | 21.668 | **0.0001** |
| | | Residuals | 28 | 1.474 | 0.564 | | |
| | | Total | 29 | 2.615 | 1.000 | | |
| **(b)** Semmeldalen (reindeer exclusion) | Grasses and rushes used as separate PFTs | Herbivory treatment | 1 | 0.273 | 0.047 | 1.893 | **0.0096** |
| | | Residuals | 38 | 5.483 | 0.953 | | |
| | | Total | 39 | 5.756 | 1.000 | | |
| | Grasses and rushes merged together as graminoids | Herbivory treatment | 1 | 0.043 | 0.011 | 0.406 | 0.376 |
| | | Residuals | 38 | 4.007 | 0.989 | | |
| | | Total | 39 | 4.049 | 1.000 | | |

(*) For the Ny-Ålesund site, PERMANOVA was first run by including all three herbivory treatments, and then separately for each pair of treatments. For the Semmeldalen site, PERMANOVA was run on both the Bray-Curtis distance matrix with grasses and rushes considered as separate PFTs and the Bray-Curtis distance matrix with grasses and rushes grouped together as graminoids.

**TABLE S6.** Results from linear mixed-effects models on the effects of herbivore removal on moss-layer depth. Parameter estimates of fixed-effects (Est.) and their 95% confidence interval (CI – lower and upper bounds) for the models on moss depth (*log-transformed*) in the long-term **(a)** goose removal experiment (Ny-Ålesund) and **(b)** reindeer removal experiment (Semmeldalen). The intercept (Int.) refers to the moss depth in the baseline herbivory treatment, whereas Est. refers to the change in moss depth in the treatment being contrasted relative to the baseline (e.g. in the contrast 'Short-excl vs Open', Int. refers to open-grazed tundra and Est. refers to the change in short-term exclosures relative to Int.). Estimates in bold indicate that their 95% CI does not include zero (i.e. statistically significant effects), whereas estimates in italic indicate that their 90% CI does not include zero (i.e. biologically interesting, close-to-significant effects). Note that (b) in Semmeldalen, no short-term exclosures were available (*Material and Methods: Experimental sites and setup*). Abbreviations: Open = open-grazed tundra; Short-excl = short-term exclosures; Long-excl = long-term exclosures.

| | Model* | Contrast | Int. | Est. | low CI | up CI | *P-value* |
|---|---|---|---|---|---|---|---|
| **(a)** Ny-Ålesund (goose exclusion) | Total moss-layer depth | Short-excl vs Open | 1.650 | **0.506** | **0.229** | **0.782** | 0.0029 |
| | | Long-excl vs Open | 1.650 | **0.707** | **0.431** | **0.983** | 0.0004 |
| | | Long-excl vs Short-excl | 2.160 | 0.202 | -0.075 | 0.478 | 0.1308 |
| | Photosynthetically-active green part | Short-excl vs Open | 0.040 | **0.711** | **0.188** | **1.235** | 0.0139 |
| | | Long-excl vs Open | 0.040 | **0.562** | **0.039** | **1.086** | 0.0383 |
| | | Long-excl vs Short-excl | 0.751 | -0.149 | -0.673 | 0.374 | 0.5294 |
| | Nearly-decomposed brown part | Short-excl vs Open | 1.390 | **0.456** | **0.156** | **0.757** | 0.0080 |
| | | Long-excl vs Open | 1.390 | **0.758** | **0.458** | **1.059** | 0.0004 |
| | | Long-excl vs Short-excl | 1.840 | **0.302** | **0.0011** | **0.602** | 0.0493 |
| **(b)** Semmeldalen (reindeer exclusion) | Total moss-layer depth | Long-excl vs Open | 0.607 | **0.502** | **0.067** | **0.936** | 0.0327 |
| | Photosynthetically-active green part | Long-excl vs Open | -0.453 | **0.586** | **0.134** | **1.040** | 0.0228 |
| | Nearly-decomposed brown part | Long-excl vs Open | 0.110 | *0.446* | *-0.067* | *0.959* | 0.0733 |

(*) Models were first run for total moss-layer depth, and then separately for the photosynthetically active green part and the nearly-decomposed brown part of the moss layer.

**TABLE S7.** Results from linear mixed-effects models on the effects of herbivore removal on vascular plant chemistry. Parameter estimates of fixed-effects (Est.) and their 95% confidence interval (CI – lower and upper bounds) for the models on nitrogen (N) concentration and carbon-to-nitrogen ratio (C:N) (both *log-transformed*) of vascular plants (five focal species) and on total live-leaf N and C pools (*log-transformed+1*) in the long-term **(a)** goose removal experiment (Ny-Ålesund) and **(b)** reindeer removal experiment (Semmeldalen). The intercept (Int.) refers to the value of the response variable in the baseline herbivory treatment, whereas Est. refers to the change in the variable in the treatment being contrasted relative to the baseline (e.g. in the contrast 'Short-excl vs Open', Int. refers to open-grazed tundra and Est. refers to the change in short-term exclosures relative to Int.). Estimates in bold indicate that their 95% CI does not include zero (i.e. statistically significant effects). Note that (b) in Semmeldalen, no short-term exclosures were available (*Material and Methods: Experimental sites and setup*). Abbreviations: Open = open-grazed tundra; Short-excl = short-term exclosures; Long-excl = long-term exclosures.

**TABLE S7.**

| | Model* | Contrast | Int. | Est. | low CI | up CI | *P-value* |
|---|---|---|---|---|---|---|---|
| **(a)** Ny-Ålesund (goose exclusion) | N concentration in *Poa arctica* (grass) | Short-excl vs Open | 1.43 | **-0.22** | **-0.38** | **-0.06** | 0.0139 |
| | | Long-excl vs Open | 1.43 | -0.01 | -0.17 | 0.15 | 0.9279 |
| | | Long-excl vs Short-excl | 1.20 | **0.21** | **0.07** | **0.36** | 0.0088 |
| | N concentration in *Equisetum arvense* (horsetail) | Short-excl vs Open | 1.42 | -0.12 | -0.29 | 0.04 | 0.1218 |
| | | Long-excl vs Open | 1.42 | -0.07 | -0.24 | 0.09 | 0.3203 |
| | | Long-excl vs Short-excl | 1.30 | 0.05 | -0.11 | 0.20 | 0.4971 |
| | C:N in *Poa arctica* | Short-excl vs Open | 2.30 | **0.24** | **0.07** | **0.41** | 0.0117 |
| | | Long-excl vs Open | 2.30 | 0.01 | -0.16 | 0.18 | 0.8519 |
| | | Long-excl vs Short-excl | 2.54 | **-0.23** | **-0.38** | **-0.08** | 0.0084 |
| | C:N in *Equisetum arvense* | Short-excl vs Open | 2.26 | 0.11 | -0.05 | 0.28 | 0.1556 |
| | | Long-excl vs Open | 2.26 | 0.07 | -0.09 | 0.24 | 0.3177 |
| | | Long-excl vs Short-excl | 2.37 | -0.04 | -0.19 | 0.12 | 0.6067 |
| | Live-leaf N pool | Short-excl vs Open | 0.12 | **0.84** | **0.44** | **1.24** | 0.0014 |
| | | Long-excl vs Open | 0.12 | **1.71** | **1.31** | **2.12** | <0.0001 |
| | | Long-excl vs Short-excl | 0.96 | **0.87** | **0.47** | **1.28** | 0.0010 |
| | Live-leaf C pool | Short-excl vs Open | 0.61 | **2.26** | **1.51** | **3.02** | 0.0001 |
| | | Long-excl vs Open | 0.61 | **3.39** | **2.63** | **4.14** | <0.0001 |
| | | Long-excl vs Short-excl | 2.87 | **1.12** | **0.36** | **1.88** | 0.0092 |
| **(b)** Semmeldalen (reindeer exclusion) | N concentration in *Bistorta vivipara* (forb) | Long-excl vs Open | 1.20 | 0.01 | -0.09 | 0.11 | 0.7977 |
| | N concentration in *Poa arctica* | Long-excl vs Open | 0.91 | 0.09 | -0.18 | 0.35 | 0.416 |
| | N concentration in *Equisetum arvense* | Long-excl vs Open | 1.40 | 0.003 | -0.14 | 0.14 | 0.9426 |
| | N concentration in *Luzula confusa* (rush) | Long-excl vs Open | 0.79 | 0.01 | -1.27 | 1.29 | 0.9388 |
| | N concentration in *Salix polaris* (dwarf-shrub) | Long-excl vs Open | 1.11 | -0.002 | -0.17 | 0.17 | 0.9704 |
| | C:N in *Bistorta vivipara* | Long-excl vs Open | 2.58 | -0.01 | -0.13 | 0.10 | 0.7654 |
| | C:N in *Poa arctica* | Long-excl vs Open | 2.82 | -0.08 | -0.36 | 0.19 | 0.4428 |
| | C:N in *Equisetum arvense* | Long-excl vs Open | 2.27 | -0.001 | -0.13 | 0.13 | 0.9825 |
| | C:N in *Luzula confusa* | Long-excl vs Open | 2.96 | 0.01 | -1.30 | 1.32 | 0.9244 |
| | C:N in *Salix polaris* | Long-excl vs Open | 2.68 | 0.01 | -0.17 | 0.18 | 0.9106 |
| | Live-leaf N pool | Long-excl vs Open | 0.80 | **0.34** | **0.09** | **0.59** | 0.0191 |
| | Live-leaf C pool | Long-excl vs Open | 2.91 | **0.47** | **0.02** | **0.93** | 0.0444 |

(*) The models on each of the five focal species refer to the N concentration and C:N of live-leaf tissue; the models on live-leaf N and C pools refer to the total N and C pools in live-leaf biomass of the five focal species, which combined made up on average 84% and 92% of the aboveground vascular plant biomass at the Ny-Ålesund and Semmeldalen sites, respectively.

**TABLE S8.** Results from linear mixed-effects models on the effects of herbivore removal on moss and soil chemistry. Parameter estimates of fixed-effects (Est.) and their 95% confidence interval (CI – lower and upper bounds) for the models on nitrogen (N) concentration and carbon-to-nitrogen ratio (C:N) (both *log-transformed*) of mosses and organic soil in the long-term **(a)** goose removal experiment (Ny-Ålesund) and **(b)** reindeer removal experiment (Semmeldalen). The intercept (Int.) refers to the value of the response variable in the baseline herbivory treatment, whereas Est. refers to the change in the variable in the treatment being contrasted relative to the baseline (e.g. in the contrast 'Short-excl vs Open', Int. refers to open-grazed tundra and Est. refers to the change in short-term exclosures relative to Int.). Estimates in bold indicate that their 95% CI does not include zero (i.e. statistically significant effects), whereas estimates in italic indicate that their 90% CI does not include zero (i.e. biologically interesting, close-to-significant effects). Note that (b) in Semmeldalen, no short-term exclosures were available (*Material and Methods: Experimental sites and setup*). Abbreviations: Open = open-grazed tundra; Short-excl = short-term exclosures; Long-excl = long-term exclosures.

**TABLE S8.**

| | Model* | Contrast | Int. | Est. | low CI | up CI | *P-value* |
|---|---|---|---|---|---|---|---|
| **(a)** Ny-Ålesund (goose exclusion) | N concentration in green mosses | Short-excl vs Open | 0.33 | *-0.15* | *-0.30* | *0.002* | 0.0523 |
| | | Long-excl vs Open | 0.33 | -0.004 | -0.18 | 0.17 | 0.9548 |
| | | Long-excl vs Short-excl | 0.17 | *0.15* | *-0.03* | *0.32* | 0.0892 |
| | N concentration in brown mosses | Short-excl vs Open | 0.02 | -0.10 | -0.30 | 0.11 | 0.2954 |
| | | Long-excl vs Open | 0.02 | -0.004 | -0.22 | 0.21 | 0.9693 |
| | | Long-excl vs Short-excl | -0.08 | 0.10 | -0.12 | 0.31 | 0.3394 |
| | N concentration in organic soil | Short-excl vs Open | -0.04 | -0.06 | -0.17 | 0.04 | 0.1891 |
| | | Long-excl vs Open | -0.04 | *-0.09* | *-0.20* | *0.01* | 0.0722 |
| | | Long-excl vs Short-excl | -0.11 | -0.03 | -0.13 | 0.07 | 0.5431 |
| | C:N in green mosses | Short-excl vs Open | 3.46 | **0.19** | **0.04** | **0.34** | 0.0172 |
| | | Long-excl vs Open | 3.46 | 0.04 | -0.13 | 0.21 | 0.5673 |
| | | Long-excl vs Short-excl | 3.65 | *-0.15* | *-0.32* | *0.02* | 0.0772 |
| | C:N in brown mosses | Short-excl vs Open | 3.73 | 0.11 | -0.08 | 0.29 | 0.2340 |
| | | Long-excl vs Open | 3.73 | 0.03 | -0.17 | 0.23 | 0.7610 |
| | | Long-excl vs Short-excl | 3.84 | -0.08 | -0.28 | 0.12 | 0.3979 |
| | C:N in organic soil | Short-excl vs Open | 3.47 | 0.05 | -0.06 | 0.16 | 0.3411 |
| | | Long-excl vs Open | 3.47 | 0.07 | -0.04 | 0.19 | 0.1575 |
| | | Long-excl vs Short-excl | 3.51 | 0.03 | -0.08 | 0.14 | 0.5990 |
| **(b)** Semmeldalen (reindeer exclusion) | N concentration in green mosses | Long-excl vs Open | -0.12 | 0.06 | -0.26 | 0.38 | 0.6241 |
| | N concentration in brown mosses | Long-excl vs Open | -0.36 | -0.0002 | -0.19 | 0.19 | 0.9974 |
| | N concentration in organic soil | Long-excl vs Open | -0.63 | 0.30 | -0.25 | 0.86 | 0.2070 |
| | C:N in green mosses | Long-excl vs Open | 3.97 | -0.07 | -0.39 | 0.26 | 0.5877 |
| | C:N in brown mosses | Long-excl vs Open | 4.10 | 0.01 | -0.20 | 0.21 | 0.9416 |
| | C:N in organic soil | Long-excl vs Open | 4.20 | -0.16 | -0.44 | 0.12 | 0.1826 |

(*) Models for moss N concentration and C:N were run separately for the photosynthetically active green part and the nearly-decomposed brown part.

**TABLE S9.** Results from linear mixed-effects models on the effects of herbivore removal on $CO_2$-fluxes. Parameter estimates of fixed-effects (Est.) and their 95% confidence interval (CI – lower and upper bounds) for the models on gross ecosystem photosynthesis (GEP), ecosystem respiration (ER) and net ecosystem exchange (NEE) in the long-term **(a)** goose removal experiment (Ny-Ålesund) and **(b)** reindeer removal experiment (Semmeldalen). The intercept (Int.) refers to $CO_2$-fluxes in the baseline herbivory treatment, whereas Est. refers to the change in $CO_2$-fluxes in the treatment being contrasted relative to the baseline (e.g. in the contrast 'Long-excl vs Open', Int. refers to open-grazed tundra and Est. refers to the change in long-term exclosures relative to Int.). Whenever a covariate (continuous predictor: PAR and soil moisture) was included in the final model, its parameter estimate and 95% CI is reported. In these cases, Est. refers to the change in $CO_2$-fluxes for each change of one unit in the covariate (see the column 'Interpretation' for further details), whereas Int. is not reported because non-biologically meaningful. Estimates in bold indicate that their 95% CI does not include zero (i.e. statistically significant effects). Note that (a) in Ny-Ålesund, no measurements of NEE were taken in short-term exclosures (*Material and Methods: Sample collection and processing*), and that (b) in Semmeldalen, no short-term exclosures were available (*Material and Methods: Experimental sites and setup*). Abbreviations: Open = open-grazed tundra; Short-excl = short-term exclosures; Long-excl = long-term exclosures.

**TABLE S9.**

| | Model | Contrasts | Interpretation | Int. | Est. | low CI | up CI | *P-value* |
|---|---|---|---|---|---|---|---|---|
| **(a)** Ny-Ålesund (goose exclusion) | Gross ecosystem photosynthesis (GEP) | Long-excl vs Open | at average PAR (594 µmol $m^{-2}$ $s^{-1}$) | -1.893 | **-3.860** | **-6.020** | **-1.700** | 0.0077 |
| | | Open × PAR | Trend in µmol $CO_2$ $m^{-2}$ $s^{-1}$ (for each 1 µmol $m^{-2}$ $s^{-1}$ increase in PAR) | | -0.001 | -0.003 | 0.001 | 0.2861 |
| | | Long-excl × PAR | | | **-0.005** | **-0.007** | **-0.003** | <0.0001 |
| | Ecosystem respiration (ER) | Short-excl vs Open | (No abiotic covariates for ER) | 1.262 | **1.200** | **0.189** | **2.200** | 0.0255 |
| | | Long-excl vs Open | | 1.262 | **2.300** | **1.294** | **3.310** | 0.0008 |
| | | Long-excl vs Short-excl | | 2.459 | **1.100** | **0.099** | **2.110** | 0.0351 |
| | Net ecosystem exchange (NEE) | Long-excl vs Open | at average PAR (593 µmol $m^{-2}$ $s^{-1}$) | -0.608 | **-1.630** | **-3.500** | **-0.242** | 0.0470 |
| | | Open × PAR | Trend in µmol $CO_2$ $m^{-2}$ $s^{-1}$ (for each 1 µmol $m^{-2}$ $s^{-1}$ increase in PAR) | | 0.0002 | -0.002 | 0.002 | 0.8497 |
| | | Long-excl × PAR | | | **-0.003** | **-0.005** | **-0.002** | 0.0003 |
| **(a)** Semmeldalen (reindeer exclusion) | Gross ecosystem photosynthesis (GEP) | Long-excl vs Open | at average PAR (481 µmol $m^{-2}$ $s^{-1}$) | -2.840 | -0.713 | -2.680 | 1.260 | 0.3722 |
| | | PAR | Trend in µmol $CO_2$ $m^{-2}$ $s^{-1}$ (for each 1 µmol $m^{-2}$ $s^{-1}$ increase in PAR) | | **-0.002** | **-0.004** | **-0.001** | 0.0055 |
| | Ecosystem respiration (ER) | Long-excl vs Open | (No abiotic covariates for ER) | 2.540 | **1.320** | **0.068** | **2.560** | 0.0429 |
| | Net ecosystem exchange (NEE) | Long-excl vs Open | at average PAR (477 µmol $m^{-2}$ $s^{-1}$) and soil moisture (44 %vol) | -0.293 | 0.609 | -1.210 | 2.430 | 0.4054 |
| | | PAR | Trend in µmol $CO_2$ $m^{-2}$ $s^{-1}$ (for each 1 µmol $m^{-2}$ $s^{-1}$ increase in PAR) | | **-0.002** | **-0.003** | **-0.0004** | 0.0126 |
| | | Soil moisture | Trend in µmol $CO_2$ $m^{-2}$ $s^{-1}$ (for each 1 %vol increase in soil moisture) | | **-0.026** | **-0.046** | **-0.006** | 0.0109 |

**TABLE S10.** Results from linear mixed-effects models on the relationships between $CO_2$-fluxes and either vascular plant biomass or moss-layer depth. Parameter estimates of fixed-effects (Est.) and their 95% confidence interval (CI – lower and upper bounds) for the models on gross ecosystem photosynthesis (GEP), ecosystem respiration (ER) and net ecosystem exchange (NEE) in the long-term **(a)** goose removal experiment (Ny-Ålesund) and **(b)** reindeer removal experiment (Semmeldalen). Est. refers to the change in $CO_2$-fluxes for each change of one unit in the predictor (see the column 'Interpretation' for further details). The intercept is not reported because non-biologically meaningful. Estimates in bold indicate that their 95% CI does not include zero (i.e. statistically significant effects).

| | Model* | Interpretation | Est. | low CI | up CI | *P-value* |
|---|---|---|---|---|---|---|
| **(a)** Ny-Ålesund (geese) | GEP ~ Live biomass | Trend in µmol $CO_2$ $m^{-2}$ $s^{-1}$ (for each 1 g $m^{-2}$ change in biomass) | **-0.023** | **-0.031** | **-0.015** | <0.0001 |
| | ER ~ Live biomass | | **0.009** | **0.004** | **0.015** | 0.0014 |
| | NEE ~ Live biomass | | **-0.012** | **-0.018** | **-0.005** | 0.0013 |
| | GEP ~ Moss-layer depth | Trend in µmol $CO_2$ $m^{-2}$ $s^{-1}$ (for each 1 cm change in moss depth) | -0.171 | -0.496 | 0.154 | 0.2839 |
| | ER ~ Moss-layer depth | | **0.152** | **0.016** | **0.288** | 0.0299 |
| | NEE ~ Moss-layer depth | | **-0.225** | **-0.381** | **-0.070** | 0.0068 |
| **(b)** Semmeldalen (reindeer) | GEP ~ Live biomass | Trend in µmol $CO_2$ $m^{-2}$ $s^{-1}$ (for each 1 g $m^{-2}$ change in biomass) | **-0.021** | **-0.035** | **-0.007** | 0.0038 |
| | ER ~ Live biomass | | **0.018** | **0.006** | **0.030** | 0.0042 |
| | NEE ~ Live biomass | | 0.001 | -0.008 | 0.009 | 0.8867 |
| | GEP ~ Moss-layer depth | Trend in µmol $CO_2$ $m^{-2}$ $s^{-1}$ (for each 1 cm change in moss depth) | **-0.783** | **-1.060** | **-0.509** | <0.0001 |
| | ER ~ Moss-layer depth | | **0.686** | **0.433** | **0.938** | <0.0001 |
| | NEE ~ Moss-layer depth | | 0.013 | -0.216 | 0.243 | 0.9062 |

(*) The predictor 'Live biomass' refers to total live aboveground vascular plant biomass; the predictor 'Moss-layer depth' refers to total moss-layer depth (including the photosynthetically active green part and the nearly-decomposed brown part of mosses). In all models, we did not include 'herbivory' as a fixed-effect as it would be highly correlated with both these predictors (cf. effects of herbivores on vascular plant biomass and moss-layer depth; Appendix 1, Tables S4 and S6).

**TABLE S11.** Results from linear mixed-effects models on the relationships between either soil temperature or soil moisture and moss-layer depth. Parameter estimates of fixed-effects (Est.) and their 95% confidence interval (CI – lower and upper bounds) for the models on soil temperature and soil moisture in the long-term **(a)** goose removal experiment (Ny-Ålesund) and **(b)** reindeer removal experiment (Semmeldalen). Est. refers to the change in either soil temperature or soil moisture for each change of one unit in moss-layer depth (see the column 'Interpretation' for further details). The intercept is not reported because non-biologically meaningful. Estimates in bold indicate that their 95% CI does not include zero (i.e. statistically significant effects).

| | **Model*** | **Interpretation** | **Est.** | **low CI** | **up CI** | ***P-value*** |
|---|---|---|---|---|---|---|
| **(a)** Ny-Ålesund (geese) | Soil temperature ~ Moss-layer depth | Trend in soil temperature (°C) (for each 1 cm change in moss depth) | 0.04 | -0.09 | 0.16 | 0.5460 |
| | Soil moisture ~ Moss-layer depth | Trend in soil moisture (%vol) (for each 1 cm change in moss depth) | -0.87 | -2.57 | 0.83 | 0.3048 |
| **(b)** Semmeldalen (reindeer) | Soil temperature ~ Moss-layer depth | Trend in soil temperature (°C) (for each 1 cm change in moss depth) | **-0.50** | **-0.67** | **-0.34** | <0.0001 |
| | Soil moisture ~ Moss-layer depth | Trend in soil moisture (%vol) (for each 1 cm change in moss depth) | 0.93 | -2.11 | 3.96 | 0.5371 |

(*) 'Moss-layer depth' refers to total moss-layer depth (including the photosynthetically active green part and the nearly-decomposed brown part of mosses).

**Supplementary Figures**

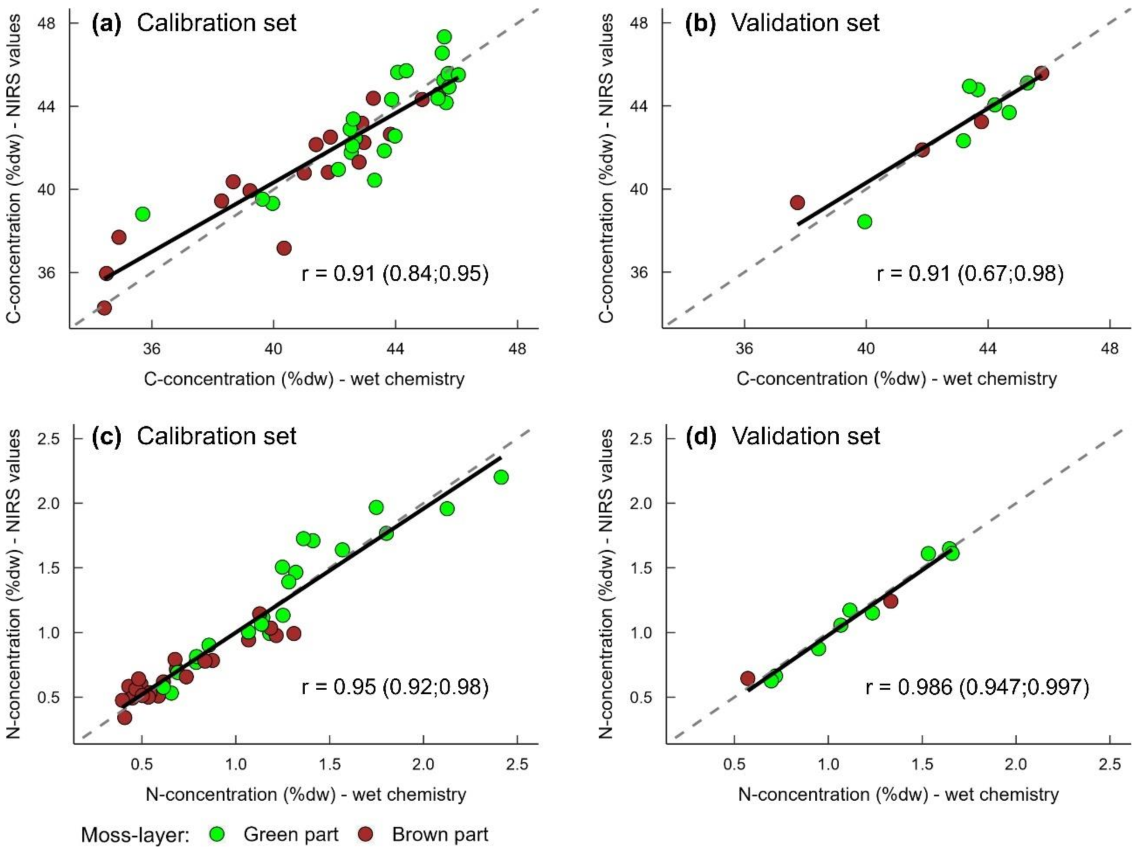


**FIGURE S1.** Correlations between moss carbon and nitrogen concentrations obtained using wet chemistry and using Near Infrared Reflectance Spectroscopy (NIRS) methodology. Correlations between values obtained with wet chemistry and values obtained with NIRS in the (a,c) calibration sample sets and (b,d) validation sample sets for carbon (C) concentration (upper panels) and nitrogen (N) concentration (lower panels) in mosses. Pearson correlation coefficients (r) and their 95% confidence interval for each relationship (solid lines) are given in each panel. The dashed lines show the ideal 1:1 relationship. Calibration models derived from (a,c) the calibration sample sets and further validated using (b,d) the validation sample sets have been used to predict C and N concentrations in moss samples (Appendix 1, Table S8).

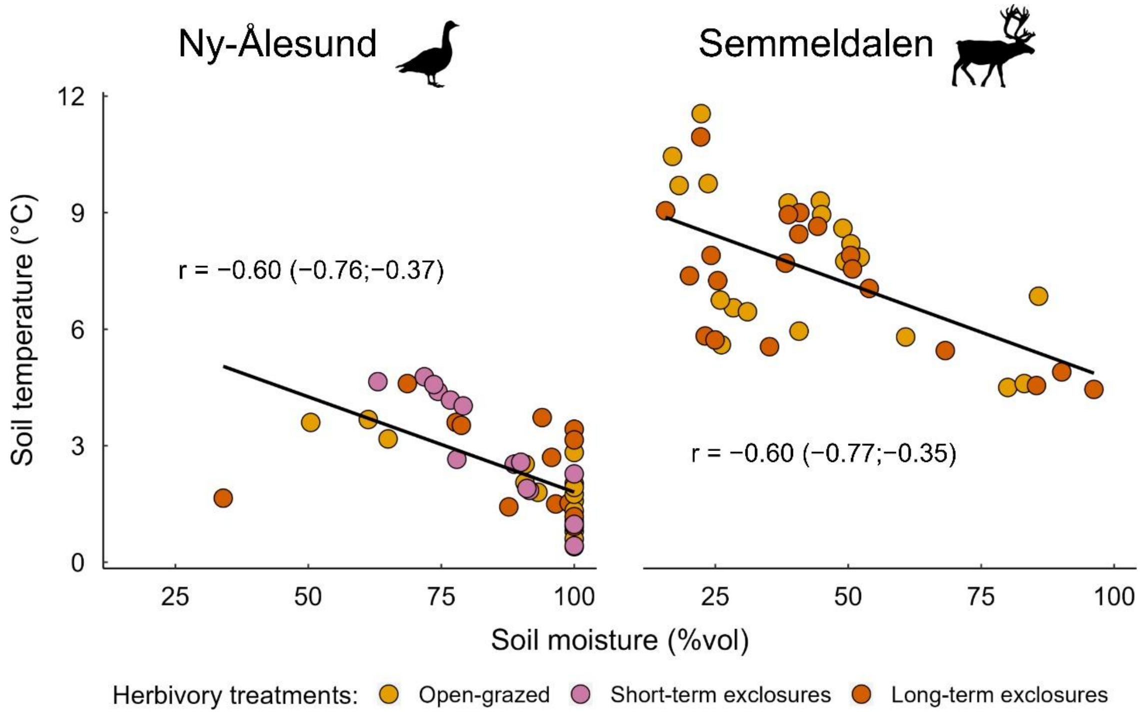


**FIGURE S2.** Correlations between soil moisture and soil temperature in Ny-Ålesund (left panel) and Semmeldalen (right panel). Pearson correlation coefficients (r) and their 95% confidence interval for each relationship (solid lines) are given in each panel.

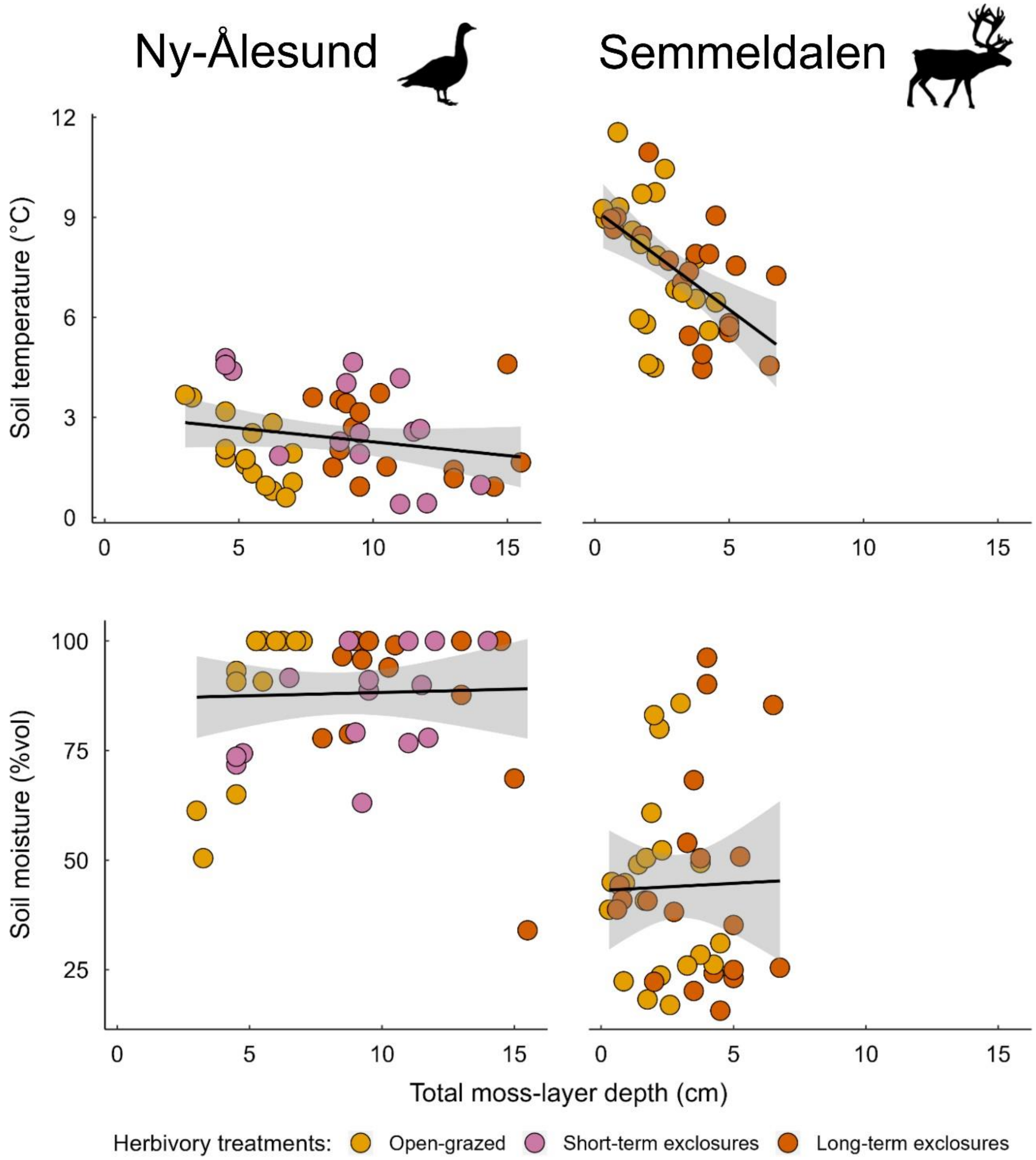


**FIGURE S3.** Relationships between either soil temperature or soil moisture and total moss-layer depth in Ny-Ålesund (left panels) and Semmeldalen (right panels). Lines and bands represent regression lines and their 95% confidence interval.
